# Conserved basal lamina proteins, laminin and nidogen, are repurposed to organize mechanosensory complexes responsible for touch sensation

**DOI:** 10.1101/2022.02.11.479800

**Authors:** Alakananda Das, Joy A. Franco, Ben Mulcahy, Lingxin Wang, Dail Chapman, Chandni Jaisinghani, Beth L. Pruitt, Mei Zhen, Miriam B. Goodman

## Abstract

The sense of touch is conferred by the conjoint function of somatosensory neurons and skin cells. These cells meet across a gap filled by a basal lamina, an ancient structure found in metazoans. Using *Caenorhabditis elegans*, we show that membrane-matrix complexes containing laminin, nidogen, and the MEC-4 mechano-electrical transduction channel reside at this interface and are instrumental for proper touch sensation. These complexes fail to coalesce in touch-insensitive extracellular matrix mutants and in dissociated neurons. MEC-4, but not laminin or nidogen, is destabilized by point mutations in the C-terminal Kunitz domain of the extracellular matrix component, MEC-1. Thus, neuron-epithelial cell interfaces are instrumental in mechanosensory complex assembly and function. Loss of nidogen reduces the density of mechanoreceptor complexes and the amplitude of the touch-evoked currents they carry. These findings imply that somatosensory neurons secrete proteins that actively repurpose the basal lamina to generate special-purpose mechanosensory complexes responsible for vibrotactile sensing.

## Introduction

Touch is our most intimate sense. It is initiated by mechanical deformation of the skin generated by direct application of static force or mechanical vibration. Among other adaptive behaviors, vibrotactile sensation enables animal communication, mate-finding, and mating (Hill, 2008). Additionally, touch sensation is instrumental for the cognitive and emotional development of humans and other primates (Cascio et al., 2019). In most animals, vibrotactile sensing is made possible by somatosensory neurons that innervate and integrate into the animal’s skin and these neurons all use mechanosensitive ion channels to convert physical deformation into electrical signals. This process of mechanotransduction is thought to occur via two, non-exclusive, biophysical mechanisms. Channels sensitive to membrane tension are thought to operate via a force-from-lipid mechanism in which sensory stimuli increase membrane tension. Other channels are proposed to operate via a force-from-filament mechanism in which physical deformation is conveyed to channels via extracellular or intracellular filaments (Goodman et al., 2023; Katta et al., 2015; Kung, 2005). Channels that use one or both mechanisms have been linked to somatosensation in vertebrates and invertebrates and osmotic sensing in microbes and plants (reviewed in Goodman et al., 2023).

Many vibrotactile sensory neurons are closely associated with epidermal cells or to specialized, epidermal-derived sensory cells, while some appear to weave paths among cells in the epidermis (Han et al., 2012; Jiang et al., 2019; Kim et al., 2012; O’Brien et al., 2012; Talagas et al., 2020; Yin et al., 2021). This association seems to emerge over time. For instance, in newly hatched *Caenorhabditis elegans* larvae, the lateral touch receptor neurons run alongside body wall muscles and these neurons become engulfed by epidermal cells during development. In adult *C. elegans* nematodes, these sensory neurons are displaced laterally such that their long, unbranched sensory neurites are positioned near the lateral midline in adults (Chalfie, 1993; Chalfie and Sulston, 1981). Lining the epidermis is a basement membrane or basal lamina (BL), a ubiquitous and conserved extracellular matrix (ECM) that surrounds tissues in all metazoans (Aumailley, 2021). Whereas the instructive role that the BL microenvironment plays in patterning somatosensory neurons within the skin is well-characterized (Yin et al., 2021), its contribution to the molecular events responsible for touch sensation remains to be fully elucidated.

Basal laminae are thin (ca. 100 nm) continuous sheets that surround several mammalian tissues (Jayadev and Sherwood, 2017; Laurie et al., 1982; Pozzi et al., 2017; Ruben and Yurchenco, 1994; Stern, 1965; Susi et al., 1967; Vracko and Strandness, 1967; Vracko, 1972, 1974; Vracko and Benditt, 1970, 1972; Younes et al., 1965). These structures are evolutionarily ancient and contain a core set of proteins conserved in all metazoans: laminin, collagen type IV, nidogen, and heparan sulfate proteoglycans or HSPS (Hynes, 2012). Laminin is a heterotrimeric protein composed of laminin-α, laminin-β, and laminin-γ subunits that typically self-assemble into continuous sheets anchored to the cell surface by a triple helical coiled-coil stalk and globular domains that bind membrane protein receptors (Cheng et al., 1997; Yurchenco and O’Rear, 1994a, 1994b; Yurchenco and Schittny, 1990; Yurchenco et al., 1986, 2002). Nidogen cross-links laminin, collagen, and the heparan sulfate proteoglycan, perlecan, in BL and other ECMs (Aumailley et al., 1993). Humans and other mammals have at least 16 distinct laminin heterotrimers (Aumailley, 2013). Conservation among vertebrate laminin isoforms and antibody cross-reactivity complicates efforts to determine which isoforms are present in specific tissues, including the skin and peripheral nervous system (Domogatskaya et al., 2012). *C. elegans*, by contrast, has only two laminin isoforms with distinct tissue localization and functions in organogenesis (Huang et al., 2003). They contain one of two laminin-α proteins (EPI-1or LAM- 3), a single laminin-β (LAM-1) protein, and a single laminin-γ (LAM-2). Although the laminins associate with the nerve ring and ventral nerve cord as well as BL surrounding other tissues, genes encoding laminins are expressed primarily outside the nervous system (Huang et al., 2003). *C. elegans* also expresses conserved laminin-binding proteins such as nidogen (NID-1), perlecan (UNC-52), syndecan (SDN-1), dystroglycan (DGN-1), and fibulin (FBL-1) (Keeley et al., 2020). These BL proteins also contribute to the organization of presynaptic structures across animal taxa [reviewed in (Sanes and Lichtman, 1999)], but their role in somatosensory function is not understood.

Genetic approaches in *C. elegans* have identified more than a dozen proteins and six touch receptor neurons (TRNs) specifically required for touch sensation (Chalfie and Au, 1989; Chalfie and Sulston, 1981). Touch evokes similar calcium transients in all six TRNs (Chatzigeorgiou et al., 2010; Nekimken et al., 2020; Suzuki et al., 2003), indicating that they all function as mechanosensory neurons. Proteins required for touch sensation include MEC-4, which is an essential pore-forming subunit of the mechano-electrical transduction (MeT) channel (O’Hagan et al., 2005), two tubulins, MEC-7 and MEC-12, and three ECM proteins, MEC-5, MEC-1, and MEC-9 (Du et al., 1996; Emtage et al., 2004). The MEC-4 channels are organized in a characteristic speckled or punctate distribution along the TRN neurites (Árnadóttir et al., 2011; Chelur et al., 2002; Chen and Chalfie, 2015; Cueva et al., 2007; Emtage et al., 2004; Katta et al., 2019; Petzold et al., 2013; Vásquez et al., 2014). The tubulins are required to form a distinctive bundle of large-diameter microtubules and for proper behavioral responses to touch (Savage et al., 1989) and to maintain proper expression of MEC-4 and several other *mec-3*- dependent genes (Bounoutas et al., 2011). Although loss of MEC-7 or MEC-12 reduces the size of touch-evoked currents, neither tubulin is required to activate MeT channels *in vivo* (Bounoutas et al., 2011; O’Hagan et al., 2005). Loss of MEC-5, MEC-1, or MEC-9 disrupts the appearance of MEC-4::YFP puncta (Emtage et al., 2004). MeT channel spacing is thought to regulate the amplitude of touch-evoked currents carried by MEC-4-dependent channels, as stronger stimuli recruit channels further away from the point of stimulus application (Katta et al., 2019). Many other proteins affect mechanosensation either directly by contributing to force transfer from the skin to the channels, or indirectly by modulating morphology, protein trafficking, and channel positioning.

Since behavioral screening methods for assessing touch sensitivity in worms require healthy, mobile animals (usually adults), mutations that are lethal or cause severe morphological defects cannot be reliably tested for defects in touch sensation. Basal lamina genes and much of the matrisome (Hynes and Naba, 2012) fall into this category. Behavioral screening methods overlook mutations that have significant effects on the structure and organization of the ECM and MeT channels and modest effects on touch response behavior. Here, we leverage new molecular tools for visualizing the well-characterized MEC-4 channel (Katta et al., 2019) and BL proteins (Keeley et al., 2020) to investigate the molecular basis of channel positioning in vibrotactile sensory neurons. Our approach combines genetic dissection with static and dynamic imaging in living animals and cultured sensory neurons, fluorescence recovery after photobleaching (FRAP), and *in vivo* electrophysiology. We show that the MEC-4 channel organizes into discrete stationary and stable clusters that co-localize with similarly stable and discrete BL puncta containing laminin and nidogen. Loss of nidogen reduces laminin and MEC-4 puncta density as well as mechanoreceptor current amplitude, confirming that nidogen contributes both the organization and function of channel-ECM complexes *in vivo*. Three ECM genes previously linked to touch sensation, *mec-5*, *mec-1*, and *mec-9*, are required for the proper organization of both the keystone MEC-4 ion channel (Du et al., 1996; Emtage et al., 2004), and the nidogen and laminin-containing BL anchor (this study). The large MEC-1 protein, which is expressed and secreted by the TRNs, is seen to play an additional role: its C-terminal Kunitz domain being required to integrate MEC-4 channels into the membrane-ECM spanning mechanosensory complex. We propose that somatosensory neurons actively repurpose conserved BL proteins to form a mechanosensory complex that bridges the neuronal plasma membrane to the extracellular matrix and is instrumental in converting touch-induced longitudinal strain into electrical signals.

## Results

The TRNs extend long (∼0.5mm), thin (∼200 nm diameter) sensory neurites that lie close to the surface of the body ensuring optimal sensitivity to indentations and are embedded within the epidermis (Figure 1A-B). A specialized extracellular matrix or ‘mantle’ (Chalfie and Sulston, 1981) fills the volume between the TRN plasma membranes and the basal membrane of the epidermis (Figure 1B), and contains proteins secreted by both the TRN and the surrounding epidermal cell. Based on overexpression of transgenes expressing fluorescent protein fusions or antibody staining, prior work has identified the collagen, MEC-5, and two Kunitz domain containing proteins, MEC-1 and MEC-9, as constituents of the ECM along TRNs (Cueva et al., 2007; Du et al., 1996; Emtage et al., 2004). MEC-1::GFP and MEC-5::CFP generated both periodic puncta and uniform fluorescence along the lateral TRNs and were proposed to co-localize based on image superposition. Although Emtage, et al (Emtage et al., 2004) reported that fluorescent protein-tagged MEC-1 and MEC-5 puncta co-localize with MEC-4 puncta, a later study using antibodies against MEC-4 and MEC-5 showed that MEC-5 does not co-localize with MEC-4 and has shorter inter-punctum intervals on average compared to MEC-4 (Cueva et al., 2007).

**Figure 1.**
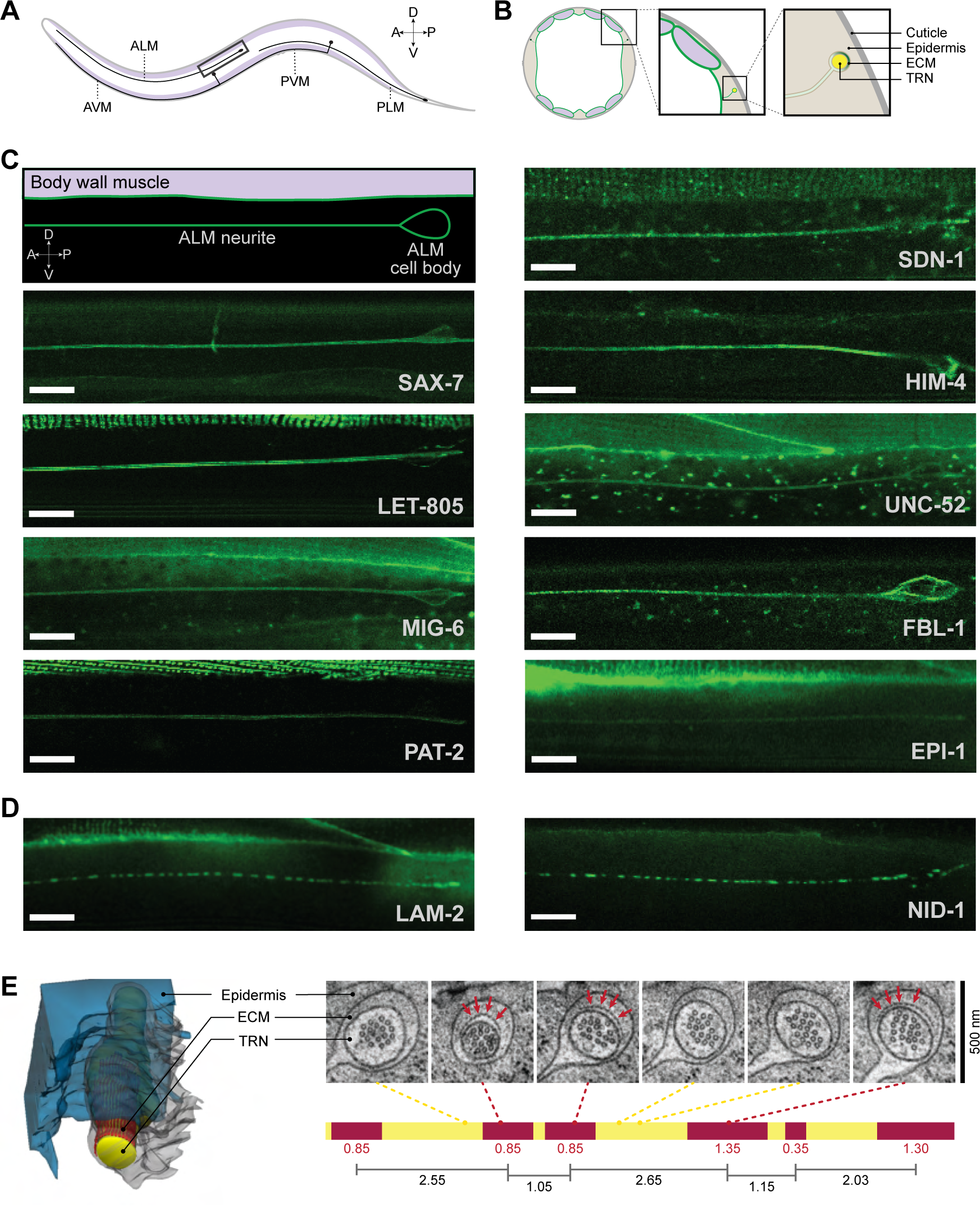
Conserved basal lamina (BL) proteins and arrays of beam-like nanostructures associate with TRNs in vivo. **A.** Schematic diagram of an adult *C. elegans* nematode, showing the positions of the TRN cell bodies and neurites relative to the body wall muscles (purple). Only the left ALM and PLM are shown. **B.** Schematic diagram of a cross-section of an adult worm, showing the body wall muscles (purple), epidermis (beige) and basal lamina (green). Enlarged view of the ALM neurons (yellow) embedded within the epidermis are shown in the middle and right panels. **C**. Nine BL proteins uniformly coat TRN neurites *in vivo.* (Top left) Schematic diagram of an enlarged view of the boxed area in panel A, showing the lateral view of an ALM cell body and part of the ALM neurite relative to the body wall muscle (purple). Representative confocal fluorescence micrographs of adult TRNs in transgenic animals expressing BL proteins visualized using CRISPR/Cas9 to insert mNG or GFP (SAX-7, LET-805) into native loci. Panels show ALM neurons coated with BL proteins, oriented as shown in the upper left panel: (left column) SAX-7 L1CAM, LET-805 myotactin, MIG-6 papillin, PAT-2 α-integrin, (right column) SDN-1 syndecan, HIM-4 hemicentin, UNC-52 perlecan, FBL-1 fibulin, and EPI-1 laminin-α. Scale bars: 10 µm. Table S1 lists the strains screened and shown in this figure. **D.** Two BL proteins localize to discrete puncta along TRN neurites. Fluorescence confocal micrographs showing LAM-2::mNG laminin-ɣ and NID-1::mNG nidogen localizing to discrete puncta distributed along the ALM neurons. Scale bar: 10 µm. **E**. (Left) A reconstruction from serial section TEM images (total length = 3 µm) of ALML in control animals [TU2769 *uIs31*[*mec-17p::GFP*] III], showing the ALM (yellow), ECM beams (red) and the plasma membrane of the engulfing epidermis (translucent blue). (Right, top) TEM micrographs of a *C. elegans* nematode (wild-type,) selected from a serial section stack, spanning approximately 10 µm. Red arrows point towards the electron dense ECM beams in the extracellular space between the ALM and the epidermis. (Right, bottom) A map of the entire serial section stack of TEM images used for the top panel, showing the relative position of the segments with (red) or without (yellow) visible ECM beams. Red numbers indicate the length (in microns) of segments with visible ECM beam arrays. Black numbers indicate the distance (in microns) between the mid-points of successive red segments. Additional examples shown in Figure S1.

Recent advances in genome editing using CRISPR/Cas9 enable engineering endogenous genetic loci in *C. elegans* to encode fusions between the encoded protein and bright, stable fluorescent proteins such as mNeonGreen (mNG) and wormScarlet (wSc) (Heppert et al., 2016; Hostettler et al., 2017; El Mouridi et al., 2017). This approach minimizes disruption to genetic regulatory elements compared to overexpression of transgenes and maximizes the ability to determine the subcellular localization of the tagged proteins and to analyze their *in vivo* dynamics compared to antibody staining. Building on a suite of transgenic *C. elegans* strains expressing mNG fusions of the *C. elegans* matrisome (Keeley et al., 2020), we built a set of dual-color transgenic strains that enabled us to visualize the BL structures at the interface between the TRNs and the ensheathing epithelial cells and their alignment with MeT channels *in vivo* and in dissociated, cultured neurons.

### ECM proteins laminin and nidogen are present in a punctate distribution along TRNs

To better understand the extracellular matrix proteins affecting touch sensation and influencing the distribution of MEC-4 channels along TRNs, we performed a visual screen to identify ECM proteins that localize to TRN neurites. We analyzed 35 *C. elegans* strains expressing fluorescent protein-labeled matrisome proteins under endogenous *cis* regulatory control at their native locus (34 strains) or within a synthetic, single copy locus (1 strain), for their localization along the TRNs. These transgenic alleles represent 27 conserved BL proteins and their putative membrane receptors (Table S1). We focused this analysis on the ALM neurons in adult worms for two main reasons: 1) no other neurons are nearby and 2) the ALM cell bodies and axons are separated from the body wall muscles and gonad, two large tissues known to express core basement membrane proteins. We found 11 proteins along the ALM neurites either in a uniform or punctate distribution (Figure 1C-D). Nine appeared to be uniformly distributed along ALM: SAX-7 L1CAM, LET-805 myotactin, MIG-6 papillin, PAT-2 α-integrin, SDN-1 syndecan, HIM-4 hemicentin, UNC-52 perlecan, FBL-1 fibulin, and EPI-1 laminin-α (Figure 1C). These observations identify three new constituents of the mechanosensory ECM (MIG-6 papillin, UNC-52 perlecan, and EPI-1 laminin-α) and reinforce prior work showing that overexpressed fusions of GFP to HIM-4 hemicentin (Vogel and Hedgecock, 2001), FBL-1 fibulin (Muriel et al., 2005), SDN-1 syndecan (Rhiner et al., 2005), SAX-7 L1-CAM (Díaz-Balzac et al., 2016) and endogenously-tagged LET-805 myotactin (Coakley et al., 2020) associate with the TRNs. With the exception of mNG::FBL-1, the introduction of the fluorescent protein encoding sequence at the endogenous locus did not significantly affect touch response behavior (Table S2). Thus, many conserved ECM proteins decorate the TRNs, suggesting that proper touch sensation depends on conserved molecular constituents of the ECM skin-neuron interface.

Two ancient and core basal lamina proteins localized to puncta distributed along TRNs: LAM-2 laminin-γ and NID-1 nidogen (Figure 1D). Given that nidogen is a laminin-binding protein (Dziadek and Timpl, 1985), the present findings are consistent with the presence of nidogen-like immunoreactivity along the ventral TRNs (Ackley et al., 2005). Notably, discrete laminin and nidogen puncta were not associated with other tissues such as the pharynx and gonad, where these proteins appear to uniformly coat tissues in continuous sheets (Clay and Sherwood, 2015; Kramer, 2005), or the in the body wall muscles in which they appear as stripes (Huang et al., 2003; Keeley et al., 2020). The TRNs have long been known to associate with a diffuse, electron-dense ECM in transmission electron micrographs (TEM) in chemically-fixed samples (Chalfie and Sulston, 1981). However, samples prepared by high-pressure freezing by us and others reveal more fine structure in the ECM. Specifically, these images contain electron-dense structures that appear as a dashed line close to the TRN plasma membrane and localize to the apical side of the TRN close to the cuticle (Frédéric et al., 2013: Figure 5A right; Hsu et al., 2014: Figure 3A; Richardson et al., 2014: figure 5J; Topalidou et al., 2012: figure 3A second from left).

To learn more about these nanoscale structures, we reconstructed them from serial-section new and published (Krieg et al., 2017) TEM data sets and found that these features consisted of arrays of beam-like structures running for nearly 1 µm parallel to the TRN plasma membrane (Figure 1E, left). On average, individual beams are approximately 20 nm wide, 8 nm thick, and separated by 8 nm and are organized in lateral arrays of 10-15 beams. They are closer to the TRN plasma membrane than they are to the epidermal cell’s basal membrane. To determine whether these structures run continuously along the length of the TRN neurite, we analyzed long serial-section datasets and found that beams were continuous in several successive sections (red segments in Figure 1E right, Figure S1A-D, Movie S1) and then disappeared (yellow segments in Figure 1E right, Figure S1A-D, Movie S1) before reappearing again. We have observed this in datasets covering at least 10 µm along the length of the three anterior TRNs from a wild-type larva (L3) and at least 8 µm stretches in the posterior TRNs from another adult wild-type animal. Notably, the ECM beams appear next to the ventral TRN, AVM, but not other adjacent neurons engulfed within the same epidermal lumen (Figure S1E). Thus, the ECM along TRNs is non-uniform along its longitudinal axis with discontinuous segments of electron dense structures and is reminiscent of the arrangement of MEC-4 channels in puncta along the ALM.

### Laminin, Nidogen, and MEC-4 co-localize to puncta that decorate TRNs

Laminin is an obligate heterotrimeric protein composed of laminin-αβγ subunits, which associate through the long coiled coil domains in each subunit. Since the laminin-αβγ subunits assemble in a 1:1:1 stoichiometry, and LAM-2 laminin-γ was detected in a punctate distribution along TRNs, we expected LAM-1 laminin-β as well as one or both laminin-α subunits, LAM-3 and EPI-1, to also localize in a similar pattern along TRNs. Homozygous LAM-1::wSc animals were sterile and showed basal lamina defects, but heterozygous animals with one tagged and one untagged copy of *lam-1* were healthy and showed punctate distribution of LAM-1::wSc along the TRNs (Figure 2A). While we detected faint EPI-1::mNG fluorescence uniformly along the ALM (Figure 1D), we did not detect any LAM-3::mNG fluorescence along the ALM. Another recent study using the same endogenously tagged LAM-3::mNG expressing strain did not detect a signal at the nerve ring (Keeley et al., 2020), although a previous study using an antibody against LAM-3 detected a strong signal at the nerve ring (Huang et al., 2003). Thus, the apparent lack of LAM-3::mNG signal along the TRNs could be due to low expression of the protein or an artifact of the position of the tag on the protein.

**Figure 2.**
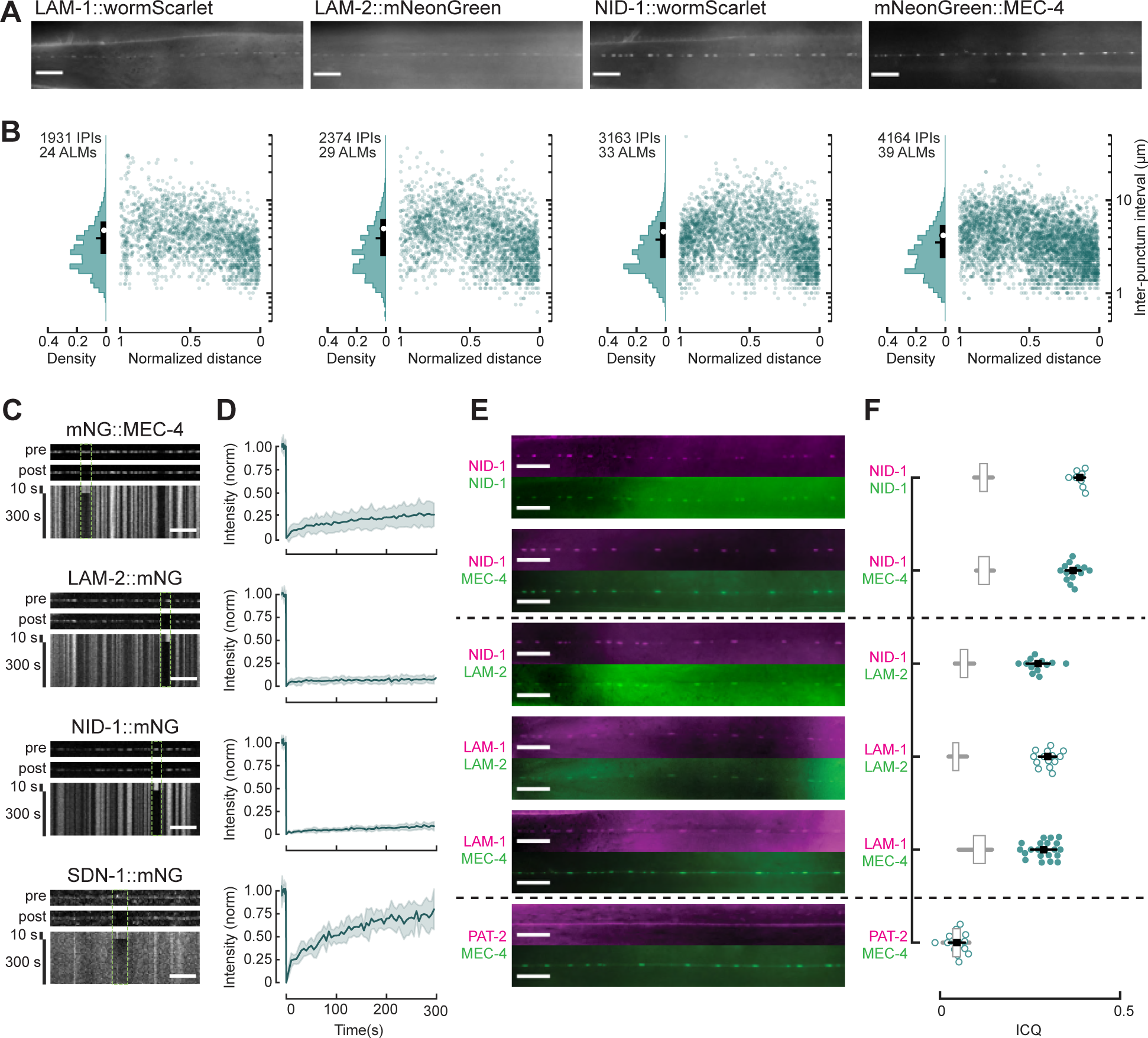
Laminin, nidogen, and MEC-4 co-localize to stationary, stable mechanosensory complexes. **A.** Representative widefield fluorescence images of puncta along ALM neurons for LAM- 1::wSc, LAM-2::mNG, NID-1::wSc and mNG::MEC-4 (from left to right). **B**. Probability density distributions plot of inter-punctum intervals (IPIs) (left) and plot of IPIs vs. normalized distance from the cell body (right; 0=closest to cell body, 1=distal tip of ALM) for each of the punctate proteins shown in panel A. White circles denote mean, horizontal black lines denote median and vertical thick black lines denote the interquartile range of the respective distributions. Mean values are collected in Table S2. One-way ANOVA comparing the IPIs of laminin, nidogen and MEC-4 showed a significant difference between the IPIs of the different proteins [*F*(3,11628) = 35.05, *p* = 1.52e-22]. Tukey HSD post hoc analysis revealed that LAM-1 IPIs were not significantly different from those of LAM-2 (*p* = 0.15) or NID-1 (*p* = 0.26) and that MEC-4 IPIs differed from all the ECM protein IPIs (*p* = 0.001). **C**. Representative, *in vivo* live imaging of (top to bottom) MEC-4, LAM-2, NID-1, and SDN-1 puncta. Top two panels show puncta before and after bleaching, respectively; the bottom panel shows kymographs (distance-time images) of a 10 s pre-bleach period and 300 s (5 minute) observation period. See also Figure S2B-E. **D.** Fluorescence recovery after photobleaching (FRAP) for MEC-4, LAM-2, NID-1, and SDN-1 puncta. Recovery was minimal for MEC-4, LAM-2, and NID-1 and substantial for SDN-1 during the observation period. Dark traces show the average trajectory of fluorescence recovery; shaded periods show the standard deviations for 23, 10, 11, and 8 neurons analyzed for MEC-4, LAM-2, NID-1, and SDN-1, respectively. One-way ANOVA comparing the fractional fluorescence recovery at 300 s showed a significant effect of the protein analyzed [*F*(3,48) = 68.73, *p* = 2.15e-17). Tukey HSD post hoc analysis revealed that the fluorescence recovery of LAM-2::mNG were not significantly different from those of NID-1::mNG (*p* = 0.9), while all other pairs were significantly different from each other. See also Figure S2B-E. **E.** Pairwise colocalization of NID-1, LAM-2, LAM-1, and MEC-tagged with wSc (magenta)and mNG (green). Fluorescence micrographs show ALM neurites in double-labeled strains. The dataset includes reference conditions for pairs expected to demonstrate high and low colocalization: High: A single protein, NID-1, is tagged with two fluorophores (NID-1::wSc, NID-1::mNG), or two subunits of an obligate heterotrimeric protein (LAM-1::wSc, LAM- 2::mNG); Low: Two TRN-expressed membrane with qualitatively distinct labeling — Uniform (PAT-2::wSc) and punctate (MEC-4::mNG). **F**. Intensity correlation quotient (ICQ) for protein pairs in panel E. Each circle (open circles: high or low reference conditions, filled circles: test strains) represents a measurement from a single ALM neurite in a dual-color strain. Black squares show the mean of the circles, and the horizontal black bars show standard deviation. Except for the PAT-2::wSc; mNG::MEC-4 pair, ICQ values computed from the same cell (circles) are significantly higher (Student’s t-test, *p* < 0.05) than that measured from simulated pairs (box plots, showing the interquartile (box) and full ranges (whiskers), excluding outliers). One-way ANOVA comparing the ICQ values across genotypes, showed significant differences in ICQ among sets of real protein pairs [*F*(5,79) = 256.73, *p* = 2.52e-47]. Horizontal dashed lines organize the strains into groups (top, middle, bottom) within which ICQs are not significantly different by Tukey post hoc analysis. Post hoc testing shows that ICQ values in both pairs in the top group differ from those for all pairs in the middle group (*p* = 0.001) and the bottom group (*p* = 0.001). Similarly, ICQ values in all pairs in the middle group differ from the pair in the bottom group (*p* = 0.001). Summary data are collected in Table S2.

The appearance of laminin-β, laminin-γ and nidogen along the TRNs shown here (Figures 1C, 2A) is similar to that reported previously for MEC-4 (Árnadóttir et al., 2011; Chelur et al., 2002; Chen and Chalfie, 2015; Cueva et al., 2007; Emtage et al., 2004; Katta et al., 2019; Petzold et al., 2013; Vásquez et al., 2014). To quantify the degree of similarity, we measured inter-punctum intervals (IPIs) for ALM-associated laminin and nidogen puncta and compared their distributions to those found for mNG::MEC-4, expressed from the single-copy *pgSi116* transgene (Katta et al., 2019). The distributions of all four proteins were comparable (Figure 2A- B), even though MEC-4 has slightly shorter IPIs on average and the distribution of IPIs of LAM-1, LAM-2, and NID-1 were more similar to one another than to mNG::MEC-4. All four proteins showed smaller IPIs closer to the cell body and larger IPIs further away (Figure 2B). Additionally, the puncta not significantly different in size — median length (μm): LAM- 1::wSc=1.29, LAM-2::mNG=1.2, NID-1::wSc=1.10 and mNG::MEC-4 1.11 (Figure S2A). Notably, these dimensions mirror the average length of the ECM beam arrays observed in ssTEM datasets (Figure 1E, Figure S1), suggesting that the protein puncta observed in living animals could be related to electron-dense structures seen in fixed samples.

Given their disparate positions within the skin-nerve complex, the MEC-4 ion channel might be expected to have distinct dynamics from the extracellular BL proteins nidogen and laminin. We tested this idea using time-lapse imaging and fluorescence recovery after photobleaching (FRAP) to determine the mobility and stability of the channel and ECM puncta. In time-lapse imaging sessions, represented as kymographs in Figure 2C, we observed that MEC-4, LAM-2 and NID-1 puncta are immobile over a time course of several minutes. During a 5-minute recovery period, photobleached MEC-4 puncta recovered only about 25% of initial fluorescence levels and photobleached LAM-2 and NID-1 puncta did not recover any detectable fluorescence (Figure 2D, Figure S2D). The limited mobility of laminin and nidogen observed in mechanosensory complexes is specific to this environment, since laminin and nidogen in the basal lamina around pharynx recover nearly 10 and 20% of their fluorescence, respectively, 5 minutes after photobleaching (Keeley et al., 2020). Additionally, the modest recovery of MEC-4 fluorescence after photobleaching is not a general property of proteins associated with the ALM plasma membrane. SDN-1 syndecan, a transmembrane heparan sulfate proteoglycan that uniformly coats the ALM (Figure 1D), recovered ∼75% of its baseline fluorescence in the same time frame (Figure 2D). These findings cannot be due to a general lack of fluidity or mobility in the TRN plasma membrane, since myristoylated-GFP expressed in the TRNs recovers more than half of its fluorescence within only 1 minute after bleaching (Figure S2B,C,E). The greater apparent mobility of MEC-4 relative to LAM-2 and NID-1 could reflect exchange of channel proteins between punctate and diffuse pools (see below). Collectively, these findings show that MEC-4, LAM-2, and NID-1 molecules form stable, largely immobile, micron-scale structures along TRN neurites *in vivo* and the unexpected similarity in their dynamics provides additional support for the idea that these proteins function together.

Based on the similar distribution and dynamics of laminin, nidogen, and MEC-4 in the ALMs (Figure 2A-D), we hypothesized that they belong to the same discrete puncta. To test this idea directly, we developed dual-color transgenic animals expressing wSc- and mNG-tagged protein pairs (Figure 2E). We quantified co-localization of protein pairs using two methods: intensity correlation quotient or ICQ (Figure 2F) and object co-localization (Figure S2F, S2G). The ICQ represents co-variation in fluorescence intensity as a single value between ™0.5 and 0.5 (Li et al., 2004), denoting perfect anticorrelation and correlation, respectively. We validated both co-localization metrics by comparing the results for real and simulated image pairs, which we generated by randomly selecting red and green channels from different cells within each dataset. As expected, ICQ values (Figure 2F, boxplots) and the fraction of co-located red and green puncta (Figure S2F-G, boxplots) were close to zero for the simulated pairs. Thus, two lines of evidence indicate that laminin and nidogen puncta strongly co-localize with each other and with MEC-4 puncta (Figures 2E-F, S2F-G).

To place these metrics in context, we also analyzed strains with red and green fluorophores either on the same protein (NID-1::wSc/NID-1::mNG) or on two proteins known to be a part of an obligate heterotrimer (LAM-1::wSc/LAM-2::mNG). The slightly lower ICQ for LAM-1/LAM-2 pair reflects the high background laminin fluorescence from surrounding tissues. Importantly, ICQ values for NID-1/MEC-4 were similar to those measured for the NID-1/NID-1 control (Figure 2F, Table S2). The ICQ for NID-1/LAM-2 and LAM-1/MEC-4 pairs were similar to each other and to the LAM-1/LAM-2 control (Figure 2F, Table S2). To determine whether the small size of TRN neurites (200-300 nm in diameter) might yield spurious intensity correlation, we also determined ICQ values for a pair of proteins that differ in their distribution with the ALM neurites. For this purpose, we chose PAT-2 α-integrin and MEC-4, because they are both membrane proteins expressed by the TRNs (Chen and Chalfie, 2014; Driscoll and Chalfie, 1991; Taylor et al., 2021) and because PAT-2 appears uniformly distributed along the ALM while MEC-4 is punctate (Figure 1C, Figure 2E). As expected for proteins that do not colocalize to common structures but are present in the same diffraction-limited volume, the PAT- 2/MEC-4 pair yields ICQ values close to zero. Unlike other protein pairs, this value was not significantly different from the ICQ calculated for simulated pairs of images.

In summary, we observed that the ECM proteins laminin and nidogen assume a punctate distribution similar to MEC-4 MeT channels along TRNs. The positions of these puncta are stable over several minutes and the molecules comprising the puncta are much less mobile than other proteins in the same environment (SDN-1::mNG, myr::GFP) or the same proteins in other environments [NID-1 and LAM-2 in pharynx (Keeley et al., 2020)]. These findings imply that BL molecules are anchored to immobile structures associated with the TRNs. Additionally, we observed that the positions of nidogen, laminin and MEC-4 channel puncta strongly correlate with each other and that the fluorescence associated with each pair of proteins is strongly correlated. Based on the evidence of similar punctate distributions, mobility, and fluorescence colocalization and intensity correlation, we propose that the matrix (laminin and nidogen) and membrane (MEC-4) components function together as mechanosensory complexes.

### MEC-4 released from mechanosensory complexes redistributes into a diffuse membrane pool

Prior evidence that loss of *mec-5*, *mec-1*, or *mec-9* function disrupts MEC-4 puncta (Emtage et al., 2004) suggested to us that laminin and nidogen puncta might also depend on *mec-5*, *mec-1,* and *mec-9*. To test this idea, we analyzed NID-1::wSc in *mec-5(u444)*, *mec-1(e1738)* and *mec–9(u437)* null mutants. NID-1::wSc puncta were not detected along any of the mutant ALM neurons, which were visualized by expression of a cytoplasmic GFP marker (Figure 3A). No defects in NID-1::wSc fluorescence were observed in the BL around the pharynx (Figure 3B), gonad and body wall muscle (Figure S3A), however. Similar results were obtained using LAM-2::mNG (Figure S3A,B). Thus, in *mec-1*, *mec-5* and *mec-9* mutants, laminin and nidogen are lost specifically along the TRNs while remaining intact in the other basement membranes.

**Figure 3.**
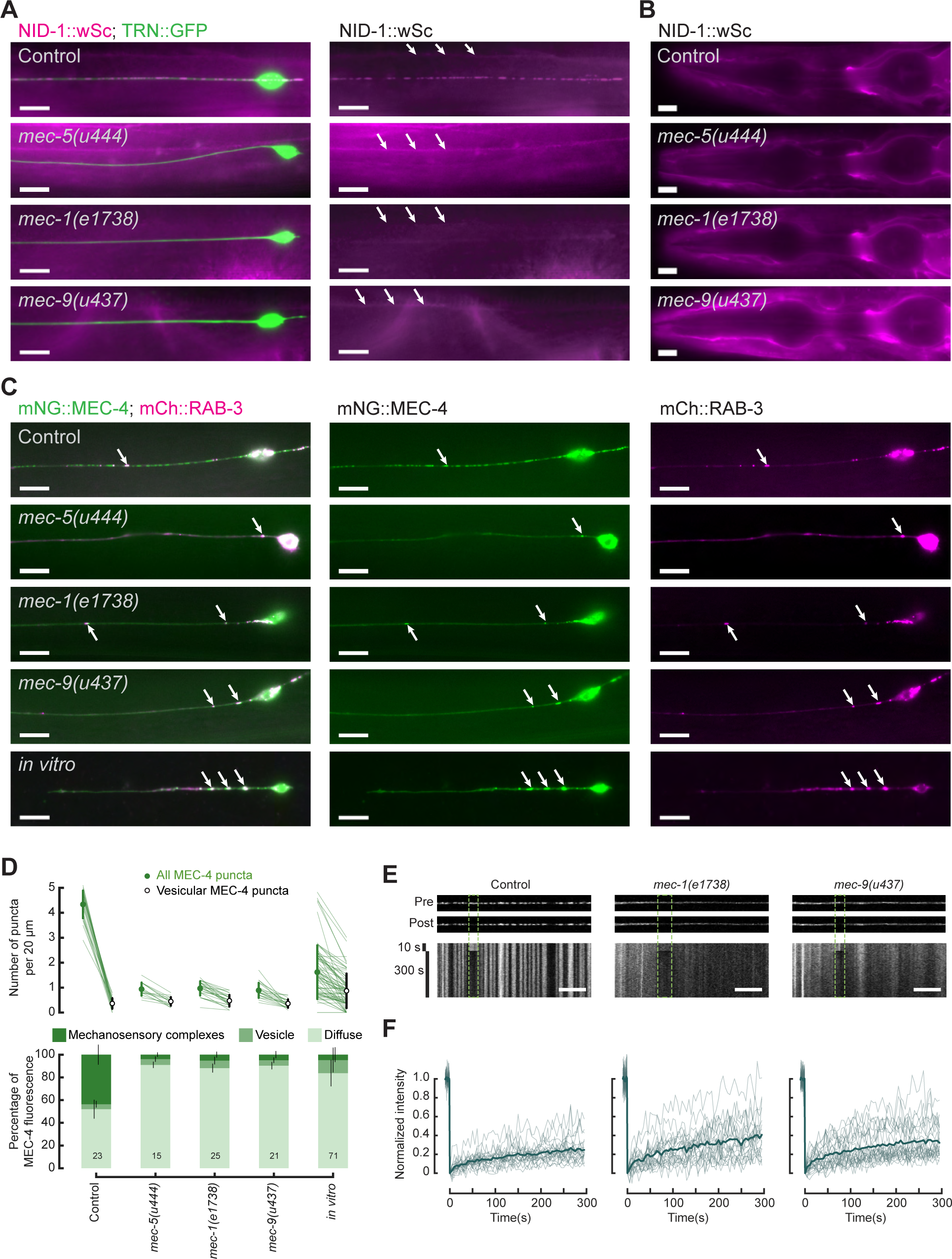
The TRN-associated extracellular matrix is essential to stabilize mechanosensory complexes in vivo and in vitro, but not for sequestering MEC-4 into RAB-3-positive vesicles. **A**. NID-1 puncta are absent in *mec-5*, *mec-1*, and *mec-9* null mutants. Representative fluorescent micrographs of animals co-expressing NID-1::wSc and a cytoplasmic GFP marker in the *mec-5(u444)*, *mec-1(e1738)* and *mec-9(u437)* mutant TRNs lack discrete NID-1 puncta in the ALM but retain fluorescence in the adjoining body wall muscle (white arrows). See also Figure S3. Scale bars are 10 µm. **B.** NID-1::wSc fluorescence in BL around the pharynx and nerve ring are not affected in *mec-5(u444)*, *mec-1(e1738)* and *mec-9(u437)* mutant animals. Scale bars are 10 µm. **C.** Representative fluorescent micrographs of TRNs co-expressing mNG::MEC-4 and mCherry::RAB-3 showing puncta co-localization. MEC-4 puncta are markedly reduced in *mec-5(u444)*, *mec-1(e1738)* and *mec-9(u437)* mutant animals and in TRNs cultured *in vitro* compared to control animals. Residual MEC-4 punctate structures often co-localize with RAB-3 (white arrows). Scale bars are 10 µm. **D**. MEC-4 puncta are significantly reduced in *mec-5*, *mec-1* and *mec-9* null mutants compared to control ALM neurons *in vivo* and dissociated, cultured TRNs *in vitro*. (Top) Plot showing the density of all detected mNG::MEC-4 puncta (green) and vesicular mNG::MEC-4 puncta (defined as puncta co-localizing with mCherry::RAB-3 puncta, white) in each cell. Pairs of data points from individual cells are connected by green lines. Circles denote mean and thick vertical lines show the standard deviation. One-way ANOVA shows significant differences in total MEC-4 puncta density [*F*(3,80) = 426.44, *p* = 4.32e-49] but no significant difference in the vesicular MEC-4 puncta density [*F*(3,80) = 1.77, *p* = 0.16] between different groups. Tukey-HSD post hoc analysis shows that total MEC-4 puncta density is significantly higher in control animals compared to other groups. See also Figure S3F. (Bottom) Stacked bar plots representing the fraction of MEC-4 in mechanosensory complexes (dark green), in RAB-3 positive vesicles (medium green) and in the diffuse pool (light green) show that MEC-4 is mostly redistributed from the mechanosensory complexes to the diffuse pool in the ECM mutants. One-way ANOVA statistics of diffuse MEC-4 vs. genotype: F(3,80)=273.57, p=6.00e-42; One-way ANOVA statistics of MEC-4 in mechanosensory complexes vs. genotype: F(3,80)=304.05, p=1.26e-43. Tukey-HSD post hoc analyses show that control animals have significantly lower percentage of MEC-4 in diffuse pool and significantly higher percentage of MEC-4 in mechanosensory complexes compared to other groups) while the vesicle pool remains relatively unaffected (One-way ANOVA statistics of vesicular MEC-4 vs. genotype: F(3,80)=2.87, p=0.04; Tukey-HSD post hoc analysis shows no significant difference between all pairs except control and *mec-1(e1738)*). The same dataset was used for analysis in both the top and bottom panels. For ALMs *in vivo*, analysis was limited to the region 0-200 µm from the cell body due to a red co-injection marker expressed in head neurons obscuring the mCherry::RAB-3 signal in TRNs. For TRNs cultured *in vitro* the entire neurite was analyzed. Note: The MEC-4 distribution in TRNs cultured *in vitro* was excluded from statistical analysis since the sample preparation, image acquisition and analysis conditions were distinct. Summarized data are collected in Table S2. **E**. Representative kymographs of MEC-4 puncta *in vivo* (left to right): control, *mec-1(e1738)*, and *mec-9(u437)*. Top two panels in each column show puncta before and after bleaching, respectively; the bottom panel shows kymographs (distance-time images) of a 10 s pre-bleach period and 300s (5 minute) observation period. Control data is replicated from Figure 2C, mNG::MEC-4. Scale bars are 10 µm. **F**. Fluorescence recovery after photobleaching (FRAP) for MEC-4::mNG in wild-type, *mec-1(e1738)*, and *mec-9(u437)*. Dark traces show the average trajectory of fluorescence recovery of MEC-4 fluorescence; light traces show the results for individual trials for 23, 22, and 25 ALMs in the indicated genotypes. Recovery is accelerated for diffuse MEC-4 fluorescence in *mec-1(e1738)* and *mec-9(u437)* relative to discrete puncta in control. To facilitate comparisons, control data is reprinted from Figure 2D, mNG::MEC-4. One-way ANOVA comparing the maximum fractional fluorescence recovery at 300 s of mNG::MEC-4 in control, *mec-1(e1738)* and *mec-9(u437)* animals showed a significant difference effect of genotype [*F*(2,67) = 3.35, *p* = 0.04]. Tukey-HSD post hoc analysis revealed that the fluorescence recovery between control and *mec-1(1738)* groups were significantly different, while there was no significant difference between other pairs of groups. See also Figure S3G.

Consistent with previous reports (Emtage et al, 2004), we observed that mNG::MEC-4 puncta were markedly reduced in *mec-5(u444)*, *mec-1(e1738)* and *mec–9(u437)* mutant animals *in vivo*. We next asked if physically separating TRNs from the native ECM would phenocopy this effect. To achieve this goal, we dissociated cells from mNG::MEC-4 transgenic embryos and cultured them on coverslips that we engineered to be covered in thin strips of adhesive proteins (STAR*Methods). This approach constrained TRNs to extend straight neurites reminiscent of their *in vivo* morphology. Even though dissociated embryonic TRNs are known to express *mec-1* and *mec-9* (Zhang et al., 2002), mNG::MEC-4 puncta were reduced *in vitro* (Figure 3C,D). The few bright punctate structures seen in mutants *in vivo* and in TRNs *in vitro* often co-localized with the vesicle marker mCherry::RAB-3 (Figure 3C,D). In contrast to stationary MEC-4 puncta in control animals, mobile MEC-4 puncta often exhibited bidirectional movement with frequent pauses and reversals, reminiscent of vesicles transporting cargo (Figure S3C-E) and were distinct from stationary MEC-4 aligned with stationary NID-1 puncta *in vivo* (Figure S3E). This suggests that punctate MEC-4 structures seen in *mec-5*, *mec-1*, *mec-9* animals *in vivo* and in cultured TRNs *in vitro* represent MEC-4 in vesicles. The average number of RAB-3 labeled puncta and the average number of MEC-4 puncta that co-localize with RAB-3 puncta does not vary significantly between control and ECM mutant genotypes *in vivo* (Figure S3F). In TRNs cultured *in vitro*, the occurrence of punctate MEC-4 and RAB-3 varied among cells, but most MEC-4 puncta co-localized with RAB-3 puncta. The vesicular MEC-4 comprises a relatively minor fraction of the total MEC-4 in ALM neurites on average (less than 10%), and this fraction also does not change appreciably between control and ECM mutants *in vivo* (Figure 3D). For simplicity, we discount the contribution of vesicular MEC-4 fluorescence in future measurements.

ECM mutants exhibit a significant increase in the fraction of mNG::MEC-4 that appears uniformly distributed along the TRN (Figure 3D). To understand whether these diffuse MEC-4 molecules are more mobile than those in discrete puncta, we performed FRAP studies of diffuse mNG::MEC-4 in mutants and compared these results to FRAP studies of discrete puncta in control animals (Figure 3E-F). This analysis focused on the *mec-1* and *mec-9* genes because they are co-expressed in the TRNs (Du et al., 1996; Emtage et al., 2004). The *mec-5* gene is not expressed in the TRNs (Du et al., 1996). As is evident from Figure 3F, FRAP profiles were more variable in *mec-1* and *mec-9* mutants than in control animals. The mutant dataset includes profiles in which the fractional recovery approached >50% after 5 minutes as well as those in which the fractional recovery was less than 20% in the same time frame. On average, both mutants had an increased fractional fluorescence recovery (*mec-1(e1738)*: 0.39±0.23 and *mec-9(u437)*: 0.34±0.17 (mean±sd)) relative to control (0.25±0.13, mean±sd). Although the difference is modest, it was statistically significant (One-way ANOVA). Post-hoc testing (Tukey HSD) revealed a statistically significant increase in the fractional fluorescence recovery at 300 s between in *mec-1(e1738)* mutants relative to control, but not between *mec-9(u437)* mutants vs. controls (Figure S3G). Notably, the MEC-4 molecules present in *mec-1* and *mec-9* mutants are much less mobile than other membrane proteins such as SDN-1 (Figure 2D), a finding which implies that other factors act in parallel to limit MEC-4 mobility. Taken together, these findings imply that the neuronal factors, *mec-1* and *mec-9*, are required to properly structure the mechanosensory BL and that, in their absence, MEC-4 is redirected into a diffuse membrane pool. This analysis also provides new insight into the role RAB-3 positive vesicles might play in the subcellular distribution of mechanosensory channels in sensory neurons *in vitro* and *in vitro*.

### Neither MeT channels nor TRN microtubules are required to assemble the matrix components of the mechanosensory complexes

Having established that MEC-4, laminin, and nidogen co-localize to stable, stationary complexes in the ALM neurons, we next used genetic dissection to learn more about how these complexes are assembled. We first measured laminin and nidogen puncta in *mec-4* null mutants. LAM-2 and NID-1 puncta were unaffected by loss of *mec-4* (Figure 4A-C, Table S2), indicating that although MEC-4 is an essential pore-forming subunit of the MeT channel in the TRNs (O’Hagan et al., 2005), it is not needed to form ECM puncta along TRN neurons. Since MEC-4 is proposed to form heterotrimers with MEC-10, DEGT-1, or both channel proteins (Árnadóttir et al., 2011; Chatzigeorgiou et al., 2010; O’Hagan et al., 2005), we asked if these other DEG/ENaC/ASIC channel subunits were required for ECM puncta formation. As found for *mec-4* single mutants, laminin puncta were unaffected in *degt-1;mec-4mec-10* triple mutants (Figure S4A-C). This finding, along with previous studies reporting that the distribution of ECM proteins MEC-1 and MEC-5 along the TRN are also not affected by loss of MEC-4 (Emtage et al., 2004), suggests that MeT channels are not sufficient by themselves to nucleate mechanosensory complexes in TRNs.

**Figure 4.**
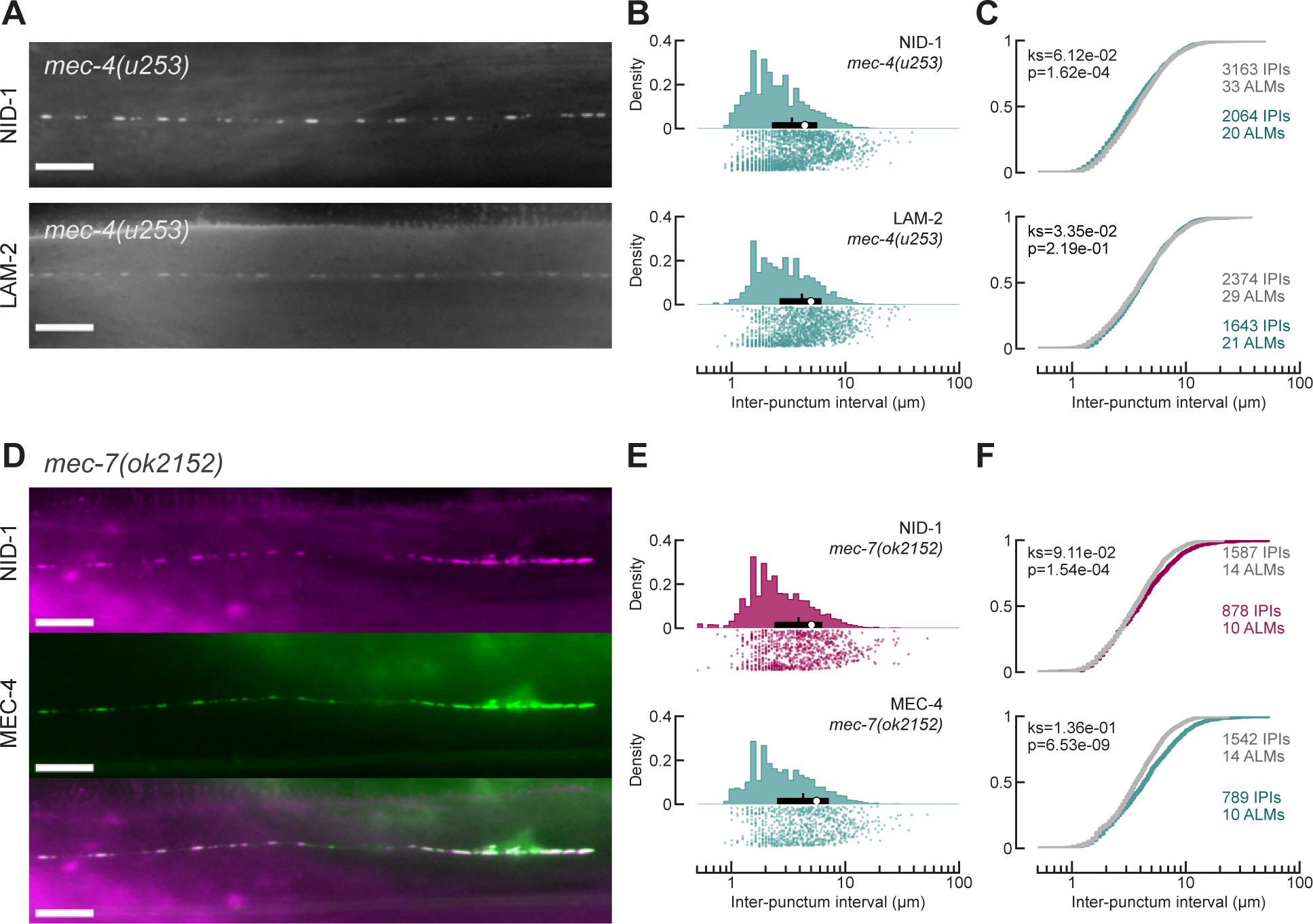
ECM mechanosensory complexes can form in the absence of MEC-4 and TRN microtubules. **A.** Representative images showing NID-1::wSc and LAM-2::mNG localize to discrete puncta distributed along the ALM neurons in *mec-4(u253)* null background. Scale bars are 10 µm. **B.** Probability density distributions plot of IPIs (top) and the individual IPIs (bottom) for NID-1 and LAM-2 in *mec-4(u253)* animals. White circles denote mean, thin vertical lines denote the median and thick horizontal lines denote the interquartile ranges of the population. Summarized data are collected in Table S2. **C.** Cumulative probability distribution of NID-1 and LAM-2 IPIs in *mec-4(u253)* animals (green lines) compared to control animals (gray lines). The results of two-sample Kolmogorov-Smirnov tests are indicated in each plot and suggest that control and mutant distributions differ for NID-1 (*p <* 0.001, top), but not LAM-2 (*p* = 0.219, bottom). See also Figure S4A-C. **D.** Representative images showing NID-1::wSc and mNG::MEC-4 puncta obtained from dual-color transgenic animals in a *mec-7(ok2152)* β-tubulin mutant. Scale bars are 10 µm. **E**. Probability density distributions plot of IPIs (top) and the individual IPIs (bottom) for NID-1 (magenta) and MEC-4 (green) puncta in *mec-7(ok2152)* animals. White circles denote mean, thin vertical lines denote the median and thick horizontal lines denote the interquartile ranges of the population. Summarized data are collected in Table S2. **F.** Cumulative distribution functions (CDF) of NID-1 and MEC-4 IPIs in *mec-7(ok2152)* animals (magenta and green lines) compared to control animals (gray lines). The results of two-sample Kolmogorov-Smirnov tests are indicated in each plot. See also Figure S4D-G.

We also found that NID-1::wSc puncta distribution along the ALM neurite was minimally affected in animals carrying a null allele of the TRN-specific β-tubulin *mec-7* gene. In the *mec-7(ok2152)* mutant background, both NID-1 and MEC-4 puncta were clearly visible in the proximal neurite (Figure 4D). Consistent with data reported earlier using *mec-7(u440)* allele (Emtage et al., 2004), MEC-4 puncta were dimmer and harder to identify in *mec-7(ok2152)* mutants, especially in the distal ALM. Although fewer MEC-4 molecules were present in the *mec-7(ok2152)* animals, the distribution between the punctate and diffuse pool was similar to that seen in control animals (Figure S4D). In contrast, NID-1 puncta were clearly visible along the entire ALM (Figure 4D). These effects are likely to account for the lower ICQ values for NID-1 and MEC-4 (Figure S4E) observed in this mutant. At the same time, the MEC-4 and NID-1 puncta that remained co-localized one another (Figure S4F-G). Thus, loss of MEC-7 β-tubulin reduces MEC-4 expression, as reported (Bounoutas et al., 2011), but does not appear to prevent formation of mechanosensory complexes.

### Mechanosensory complex formation depends on nidogen and laminin

Next, we sought to determine whether laminin and nidogen were essential to form mechanosensory complexes and to position MEC-4 channels. We could not analyze mechanosensory puncta in laminin null mutant adults because loss of any of the four laminin genes, *epi-1, lam-3, lam-1,* and *lam-2,* causes lethality in *C. elegans* embryos or larvae (Kao et al., 2006). We could, however, analyze the laminin-ɑ *lam-3(ok2030)* mutant animals that do not develop past L1 stage and small number of *epi–1(gm57)* mutants that survive to adulthood (Forrester et al., 1998). Given that laminin-ɑ is essential for laminin trimer assembly (Kao et al., 2006; Yurchenco et al., 1997), investigating MEC-4 channel position in *epi-1* and *lam-3* laminin-ɑ mutants provides insight on the role that laminin trimers play in organizing mechanosensory puncta.

We analyzed and compared MEC-4 puncta in control, *epi-1(gm57)*, and *lam-3(ok2030)* mutant L1 larvae. As in adults, MEC-4 localized to discrete puncta in control L1 larvae (Figure 5A). *epi-1(gm57)* L1 animals had MEC-4 puncta similar to control animals, with IPIs that are only marginally higher on average from control (Figure 5A-C). In the few *epi-1(gm57)* mutant animals that survive to adulthood, the ALM cell bodies and sensory neurites displayed ectopic branching and multiple segments were seen adjacent to the body wall muscle, displaced from their control position (Figure S5A). TRN neurite segments that were adjacent to the body wall muscle in *epi-1(gm57)* mutants displayed only diffuse mNG::MEC-4 fluorescence. By contrast, segments that retained their wild-type position retained discrete MEC-4 puncta with IPIs indistinguishable from control animals (Figure S5B-C, Table S2). Thus, wild-type EPI-1 laminin-ɑ is required to position the ALM neurons in the worm’s body, but dispensable for positioning mechanosensory complexes along properly attached ALM neurons. In *lam–3(ok2030)* larvae, MEC-4 puncta were either undetectable or when visible (Figure 5A), they were fewer in number and more widely separated. The increase in the reported mean and median IPI values in *lam-3(ok2030)* larvae compared to control (Figure 5B-C, Table S2) underestimates the effect of *lam-3* loss-of-function, since in 50% (17/34) of the *lam-3(ok2030)* L1 larva we imaged mNG::MEC-4 signal in the ALM could not be visualized distinctly and were thus not analyzed. Thus, while EPI-1 adheres to TRNs (Figure 1D) and is needed to maintain proper ensheathment within epidermal cells, LAM-3 is needed for mechanosensory complex assembly. The two *C. elegans* laminins proteins play distinct, but overlapping roles in mechanosensation, as they do in embryonic development (Huang et al., 2003).

**Figure 5.**
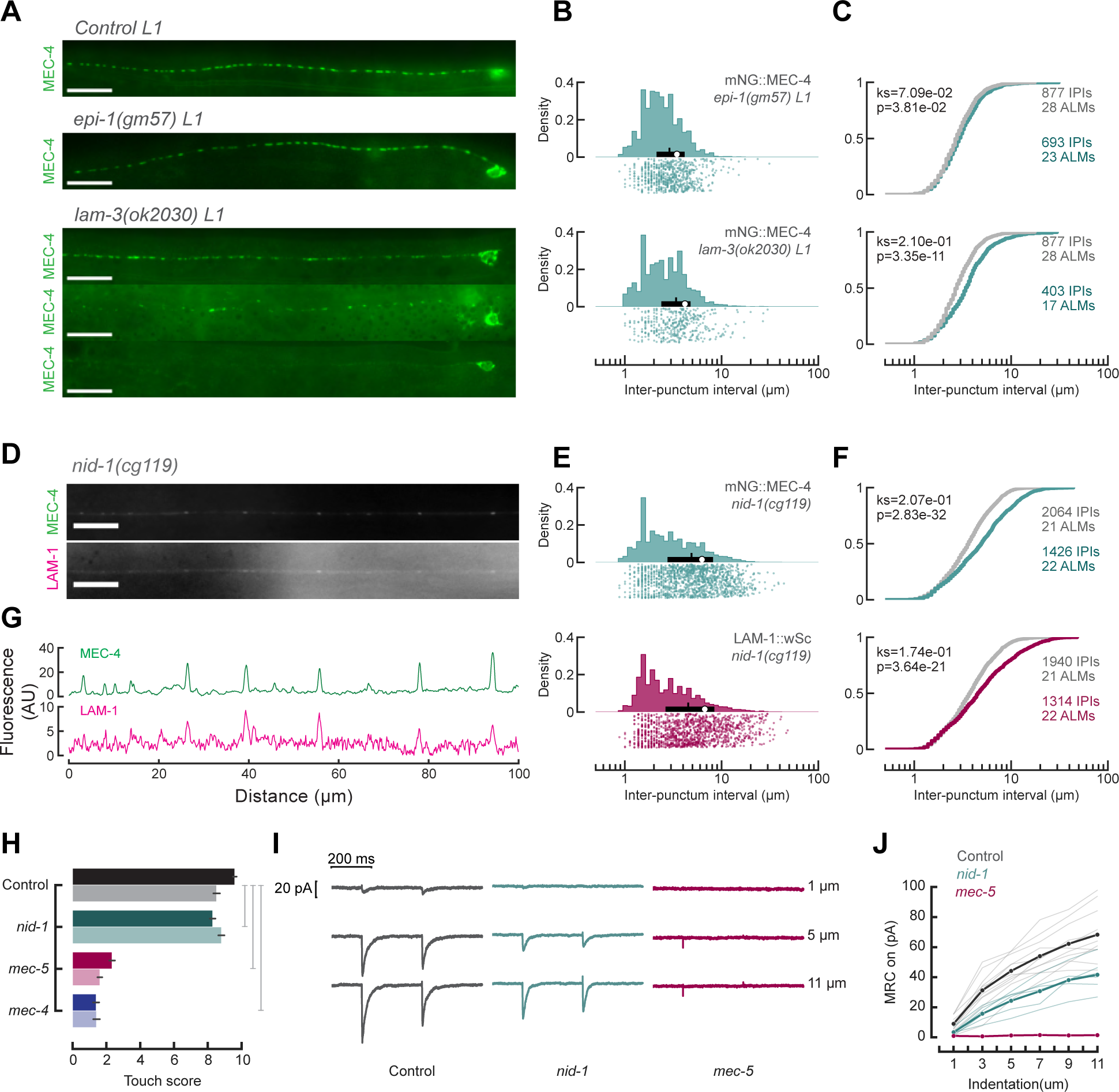
Loss of lam-3 and nid-1, but not epi-1 decreases the density of mechanosensory complexes. **A.** Representative micrographs showing mNG::MEC-4 puncta distributed along the ALM neurons in control and *epi-1(gm57)* L1 larvae, but disorganized or absent in *lam-3(ok2030)* L1 larvae. Scale bars are 10 µm. See Figure S5A for images of TRN attachment defects in *epi-1(gm57)* adult animals. **B.** Probability density distributions plot of IPIs (top) and the individual IPIs (bottom) for mNG::MEC-4 in *epi-1(gm57)* and *lam-3(ok2030)* L1 larva animals. Due to the disrupted or lack of MEC-4::mNG signal in many *lam-3(ok2030)* L1 larva animals, IPIs could be computed for only 17 out of 34 images collected. White circles denote mean, thin vertical lines denote the median and thick horizontal lines denote the interquartile ranges of the population. Summarized data are collected in Table S2. **C.** Cumulative distribution functions (CDFs) for mNG::MEC-4 IPIs in *epi-1(gm57)* and *lam-3(ok2030)* (green lines) L1 larvae compared to control L1 larvae (gray lines). The results of two-sample Kolmogorov-Smirnov tests are indicated in each plot. **D.** Representative images of mNG::MEC-4 and LAM-1::wSc obtained from dual-color transgenic animals in a *nid-1* mutant background showing dim and sparse puncta in both channels. Scale bars are 10 µm. **E**. Probability density distributions plot of IPIs (top) and the individual IPIs (bottom) for mNG::MEC-4 (green) and LAM-1::wSc (magenta) in *nid-1(cg119)* mutant animals. White circles denote mean, thin vertical lines denote the median and thick horizontal lines denote the interquartile ranges of the population. Summarized data are collected in Table S2. **F.** Cumulative distribution functions (CDFs) of MEC-4 (top) and LAM-1 (bottom) IPIs in ALM neurites. Larger intervals dominate the distribution in *nid-1* mutants (magenta and green) compared to control (gray). The results of two-sample Kolmogorov-Smirnov tests comparing CDFs are indicated in each plot. **G.** Intensity line scans derived from images in panel D showing the spatial coincidence of peak intensities in each channel. See also Figure S5D-G. **H-J.** *nid-1* loss of function decreases behavioral and neural responses to touch. (H) Touch response as a function of genotype, using wild-type (gray) and *mec-4(u253)* (blue) null animals as references show very small touch defects in *nid-1(cg119)* (teal) animals but severe touch defects in *mec-5(u444)* (magenta) animals. For each genotype, strains with or without the TRN marker *uIs31* are represented by light and dark colors respectively. Bars are the average (± SEM) response to 10 touches of *n=75* animals tested blind to genotype in 3 independent cohorts. 2-way ANOVA shows significant differences in touch-response between genotypes [*F*(3,594)=1135.12, p=1.9e-245] as well as presence or absence of the *uIs31* reporter [*F*(1,594)=6.59, p=1.05e-02]. Tukey-HSD post hoc analysis shows significant differences between all mutant genotypes compared to control (gray bars). Summarized data are collected in Table S2. (I) Mechanoreceptor currents (MRCs) evoked by indentation pulses of the indicated amplitude. Traces for control (dark gray), *nid-1(cg119)* (green) and *mec-5(u444)* (magenta) mutants are shown for a small (1 μm), medium (5 μm) or large (11 μm) indentations. Similar results obtained in 11, 6 and 2 recordings for control, *nid-1* and *mec-5* animals, respectively. (J) Peak MRC amplitude at the onset of stimulation as a function of stimulus displacement. Dark thick lines are the average of the lighter thin lines for control (dark gray), *nid-1(cg119)* (green) and *mec-5(u444)* (magenta) animals. One-way ANOVA shows significant differences in the peak current amplitude at 11 μm indentation between genotypes (F(2,16)=19.39, p=5.3e-05). Tukey-HSD post hoc analysis shows significant differences between all groups. See also Figure S5H-I.

Like laminin, nidogen is a core BL protein that is conserved in all vertebrates and invertebrates (Mayer et al., 1998). Unlike laminin, nidogen is not needed for BL assembly (Ackley et al., 2003; Bader et al., 2005; Böse et al., 2006; Dai et al., 2018; Dong et al., 2002; Kang and Kramer, 2000; Kim and Wadsworth, 2000; Murshed et al., 2000; Schymeinsky et al., 2002; Wolfstetter et al., 2019) and nidogen null mutants develop into grossly normal adult worms (Kang and Kramer, 2000). Both MEC-4 channel puncta and LAM-1 puncta were dimmer and spaced further apart in *nid-1* mutants compared to control animals (Figure 5D-F). The decrease in mechanosensory complex density was accompanied by an increase in the proportion of MEC-4 protein present in the diffuse membrane pool (Figure S5D), resulting in a low LAM- 1/MEC-4 ICQ value in the *nid-1* mutant (Figure S5E). However, a majority of the remaining MEC-4 and laminin puncta were still co-localized (Figure 5G, Figure S5F-G). Collectively, these findings reinforce the idea that MEC-4 channels redistribute to a diffuse membrane pool following perturbations of mechanosensory complexes and that BL play an instructive role in the assembly or stability of sensory complexes.

Despite the significantly lower density of mechanosensory complexes along the TRNs, *nid-1* animals were not significantly less touch sensitive. In the standard touch response assay, where the animals were stroked with an eyebrow hair, the *nid-1* animals scored only slightly lower than control animals and significantly more than the *mec-4(u253)* and *mec-5(u444)* animals (Figure 5H). Given that the force delivered during these assays far exceeds saturation (Nekimken et al., 2017), this assay is not able to detect small changes in touch sensitivity or in activation of MeT channels. Accordingly, we used *in vivo* whole-cell patch recording to directly determine how loss of *nid-1* affects the response to body indentation. To optimize our ability to detect a partial loss-of-function, we then used an ultrafast mechanical stimulator that evokes currents that are approximately five-fold larger than those evoked by slower stimuli (Katta et al., 2019). Using this approach, we found that mechanoreceptor currents or MRCs were qualitatively similar in *nid-1* mutants and wild-type controls (Figure 5I), but that MRC amplitude was decreased in *nid-1* mutants (Figure 5J). The effect of *nid-1* is specific to MRCs, since voltage-gated currents were similar to wild-type controls in all respects (Figure S5H-I). Thus, we conclude that the lower MRCs found in *nid-1* mutants arise from the decreased density of sensory complexes. This finding is consistent with computational studies indicating that stimulus intensity, speed, and channel density govern MRC amplitude (Katta et al., 2019).

### Nidogen and laminin inclusion in mechanosensory complexes is independent of direct interaction between nidogen and laminin

Wild-type, NID-1 protein contains three conserved globular domains, G1, G2 and G3 (Figure 6A). The N-terminal G1 and G2 domains are connected by a flexible linker, whereas the G2 and G3 domains are connected by a more rigid rod-like domain consisting of a series of EGF motifs (Fox et al., 1991; Patel et al., 2014). Multiple studies with recombinant mouse nidogen fragments show that perlecan binds exclusively to the G2 domain, laminin-ɣ1 binds exclusively to the G3 domain, collagen IV and fibulin-1C bind to the G2 and G3 domains, and fibulin-2 binds to all three globular domains (Fox et al., 1991; Kvansakul et al., 2001; Reinhardt et al., 1993; Ries et al., 2001). To understand which of the globular domains are important for the formation of mechanosensory complexes in worms, we first looked at the *nid-1(cg118)* allele, which encodes a large in-frame deletion of G1, G2 and part of the first EGF motif of the rod domain (Figure 6A). In *nid-1(cg118)* mutants, MEC-4 puncta distributions were similar to wild-type (Figure 6C-E top, Table S2). Thus, neither the G1 nor the G2 domains of nidogen are required to assemble or position mechanosensory complexes. The G1 and G2 domains of nidogen are likewise dispensable for expression of NID-1 protein during embryonic development and in BL associated with muscles, the somatic gonad, and neuronal tracts (Kang and Kramer, 2000).

**Figure 6.**
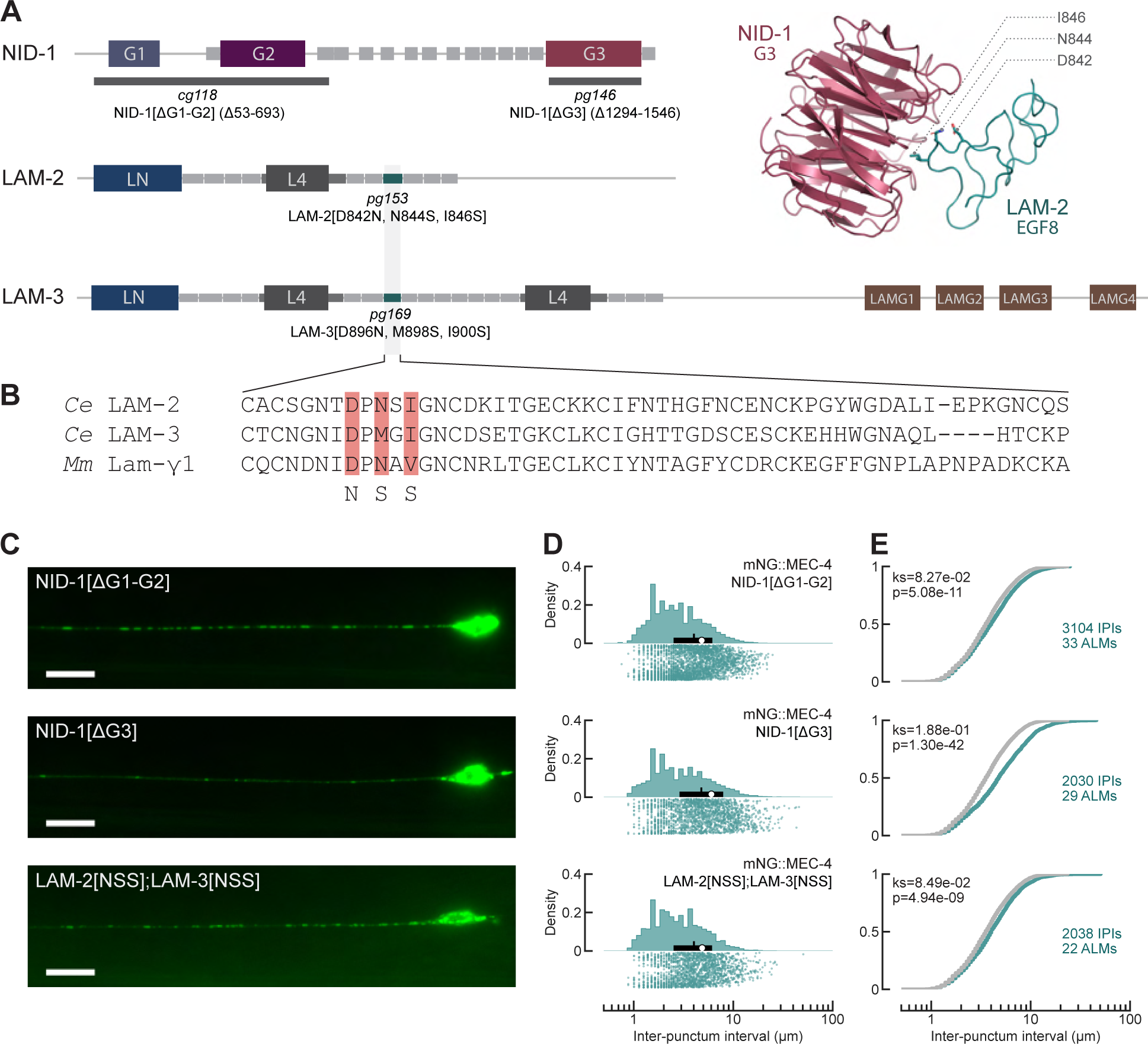
NID-1 G3 domain, but not its binding to LAM-2 or LAM-3 EGF domains is necessary for MEC-4 puncta distribution. **A**. (Left) Schematic diagram of NID-1, LAM-2 and LAM-3 proteins mapping the ΔG1-G2 (*cg118)* and ΔG3 (*pg146)* in-frame deletions in NID-1 and the point mutations in the 8th EGF domains of LAM-2 (*pg153*) and LAM-3 (*pg169*). Large boxes show the indicated globular domains. Small gray boxes show EGF domains. (Right) Homology model of *C. elegans* NID-1 G3 domain bound to LAM-2 EGF8, based on crystal structure of mouse nidogen G3 domain bound to mouse laminin-γ1 (PDB ID: 1NPE). **B.** Sequence of the LAM-2 and LAM-3 EGF8 domains predicted to bind NID-1, and its sequence alignment with its counterpart in mouse laminin-γ1. The residues mutated in the *pg153* LAM-2[NSS] and *pg169* LAM-3[NSS] alleles are highlighted in pink. **C.** Representative images of mNG::MEC-4 puncta present in *nid-1* mutant ALM neurons (top to bottom): NID-1[ΔG1-G2] (*cg118*), NID-1[ΔG3] (*pg146*), and LAM-2[NSS];LAM-3[NSS] (*pg153;pg169*). Anterior is to the left; scale bars are 10 µm. See also Figure S6. **D**. Probability density distribution plots of IPIs (top) and the individual IPIs (bottom) for mNG::MEC-4 (green) in NID-1[ΔG1-G2], NID-1[ΔG3], and LAM-2[NSS];LAM-3[NSS] mutant animals. White circles denote mean, thin vertical lines denote the median and thick horizontal lines denote the interquartile ranges of the population. Summarized data are collected in Table S2. **E**. Cumulative distribution functions (CDFs) of mNG::MEC-4 IPIs in the indicated alleles (green) vs control worms (gray, replotted from Figure 2B). The results of two-sample Kolmogorov-Smirnov tests are indicated in each plot.

Next, we examined the role of the nidogen G3 domain thought to bind to laminin-ɣ, collagen IV and fibulin (Aumailley et al., 1993; Fox et al., 1991; Mann et al., 1988, 1989; Mayer et al., 1998; Patel et al., 2014; Ries et al., 2001; Takagi et al., 2003). We used CRISPR mediated genome editing to create a NID-1[ΔG3] mutant (*pg146)* deleting the amino acids 1294-1546.

Like *nid-1(cg119)* null mutants (Ackley et al., 2003; Kang and Kramer, 2000), the NID-1[ΔG3] mutants developed into grossly wild-type adult animals. Also similar to *nid-1(cg119)*, MEC-4 puncta were dimmer and spaced further apart in NID-1[ΔG3] mutants (Figure 6C-E middle, Table S2), with a corresponding increase in the fraction of MEC-4 in the diffuse pool (Figure S6A), and a slightly reduced touch response behavior compared to control animals (Figure S6B). Thus, the NID-1 G3 domain is required for MEC-4 puncta distribution. Given that the NID- 1[ΔG3] mutant phenocopies the *nid-1(cg119)* null allele with respect to MEC-4 puncta distribution, we asked whether this is due to the inability of the truncated NID-1[ΔG3] protein to incorporate into mechanosensory complexes by visualizing a wSc-tagged NID-1[ΔG3] isoform. Consistent with the idea that the G3 domain is required to incorporate NID-1 into BL, NID- 1[ΔG3]::wSc fluorescence was severely reduced in the BL around the pharynx, body wall muscle and gonad. However, NID-1[ΔG3]::wSc was visible in neuronal-associated BL, including the nerve ring and in puncta along ALM neurons (Figure S6C-E). These findings imply that although the nidogen G3 domain is required to incorporate nidogen into conventional BL and is required for wild type mechanosensory puncta distribution, it is dispensable for inclusion into neuronal BL and into the residual mechanosensory puncta along TRNs.

Since the nidogen G3 domain has been shown to bind laminin-ɣ, collagen IV and fibulin (Aumailley et al., 1993; Fox et al., 1991; Mann et al., 1988, 1989; Mayer et al., 1998; Patel et al., 2014; Reinhardt et al., 1993; Ries et al., 2001; Takagi et al., 2003), we then asked which of these interactions are important for mechanosensory complex formation. Structural and biochemical evidence mapped the nidogen binding site on mouse laminin-ɣ1 to three critical amino acids (D800, N802, V804) on a unique loop in the 8th EGF motif (Pöschl et al., 1994; Pöschl et al., 1996; Takagi et al., 2003) (Figure 6B). Each of these residues when individually mutated reduced the binding affinity between nidogen and laminin by several orders of magnitude (Pöschl et al., 1994). In a proteome-wide search for a similar motif in *C. elegans*, we found that the 8th EGF motif on both laminin-ɣ LAM-2 and laminin-α LAM-3 contained this loop with partially conserved residues as shown in Figure 6A-B. We predicted that the EPI-1 containing laminin has only one binding site for nidogen (on LAM-2) while the LAM-3 containing laminin has two binding sites for nidogen (on both LAM-2 and LAM-3). We sought to disrupt the nidogen-laminin binding interface by engineering mutations in the putative nidogen binding residues in both LAM-2 (LAM-2[D842N, N844S, I846S]) and LAM-3 (LAM-3[D896N, M898S, I900S]). We refer to these triple amino acid substituted alleles as LAM-2[NSS] and LAM-3[NSS], respectively. We chose these amino acid substitutions because they had the largest fold-change in binding affinity between mammalian nidogen and laminin-γ1 fragments *in vitro* (Mayer et al., 1998; Pöschl et al., 1996).

Unlike *lam-2* and *lam-3* null mutants, LAM-2[NSS], LAM-3[NSS] as well as the LAM-2[NSS];LAM-3[NSS] double mutant animals were viable with no apparent defects in gross morphology or development. LAM-2[NSS]::mNG showed a similar distribution as WT LAM-2::mNG in all tissues (Figure S6C-E). As expected for loss of nidogen-laminin binding, NID-1::wSc fluorescence was absent in the BL lining the pharynx and body wall muscle in LAM-2[NSS];LAM-3[NSS] double mutants (Figure S6C-E). NID-1::wSc fluorescence was reduced but not completely lost around the same tissues in the LAM-2[NSS] single mutant (Figure S6C-E), indicating that the nidogen binding site on LAM-3 was sufficient for recruiting nidogen to BL with LAM-3 containing laminin. Conversely, NID-1::wSc fluorescence was similar to the wild-type control in the LAM-3[NSS] single mutant (Figure S6C-E), indicating that the nidogen binding site on LAM-2 which is present in both types of laminin is also sufficient for recruiting nidogen. Despite these dramatic effects in conventional basal lamina, LAM-2[NSS];LAM-3[NSS] double mutants appeared to retain NID-1 in the nerve ring (Figure S6D) and also retained a near wild-type puncta distribution of MEC-4, NID-1 and LAM-2 along the ALM (Figure 6C-E bottom, Table S2, Figure S6E). Collectively, these findings suggest that 1) nidogen redundantly binds to LAM-2 and LAM-3 using similar binding interfaces, 2) laminin-nidogen binding is not essential for the recruitment of either laminin, nidogen or MEC-4 into mechanosensory complexes, 3) the nidogen G3 domain likely interacts with another factor that regulates the spacing of the mechanosensory complexes along TRNs.

The other likely nidogen G3-binding factors are collagen IV or fibulin. Cross-linking of laminin and collagen networks via nidogen have been shown to be important for the integrity of many types of BL (Hamill et al., 2009). *C. elegans* expresses two canonical collagen IV proteins, EMB-9 and LET-2, neither of which have been detected along TRNs in our screen (Table S1) or in previous studies (Graham et al., 1997). The only known collagen-like protein known to be present around TRNs is MEC-5, which is significantly smaller in length than the canonical collagen IV proteins. It is well-known that loss of MEC-5 disrupts touch sensation (Du et al., 1996; Gu et al., 1996) and we have shown in Figure 3 that mNG::MEC-4 and NID-1::wSc puncta along the ALM are lost in the *mec-5* mutant animals. However, since the specific binding sites between collagen IV and nidogen are unknown and we have not been able to fluorescently label MEC-5 without affecting its function, we have not been able to test whether NID-1’s collagen IV binding activity is necessary for mechanosensory complex formation. *C. elegans* has one fibulin encoding gene, *fbl-1*, and fibulin protein is present along the ALMs (Figure 1C, and (Muriel et al., 2005)). The mNG::FBL-1 strain was touch insensitive (Table S2), but *fbl-1* loss of function mutants retained an essentially wild-type distribution of MEC-4 puncta along the ALM neuron (Table S2). Further experiments are necessary to fully understand the role of nidogen binding to collagen and fibulin in mechanosensory complex formation and function.

### MEC-1 connects the MEC-4 channel to ECM components of the sensory complex

The *mec-1* gene is predicted to encode a large protein containing fifteen Kunitz (Ku) domains [Figure 7A and (Emtage et al., 2004)] and similar motifs are found in peptide toxins that bind to ion channels including ASIC channels (Mishra, 2020; Cristofori-Armstrong and Rash, 2017). This raises the possibility that these toxins mimic endogenous Ku domain proteins that bind to the extracellular domains of DEG/ENaC/ASIC channels. Consistent with this idea, MEC-1 Ku15 shares 33% sequence identity with the snake neurotoxin MitTx-α (Bohlen et al., 2011) (Figure 7B). We investigated this idea *in vivo*, leveraging known and new missense mutations affecting MEC-1 Ku15. The existing *mec-1(e1526)* and *mec-1(u811)* alleles lead to the expression of full-length MEC-1[R1804C] and MEC-1[C1808Y], respectively. Unlike the residue affected in *e1526* allele, the one affected in *u811* is conserved and predicted to form a disulfide bond likely to be integral to folding Ku15 (Figure 7B). *mec-1(e1526)* and *mec-1(u811)* animals are known to be touch insensitive (Emtage et al., 2004), but how these mutations affect all elements of mechanosensory complexes is not known. Using CRISPR/Cas9 genome editing, we mutated C1808 of Ku15 tyrosine (encoded by the *pg164* allele), to recreate the *u811* mutation. Using dual-color transgenic animals expressing NID-1::wSc and mNG::MEC-4, we found that both *e1526* and *pg164* alleles retained nidogen puncta but very few mNG::MEC-4 puncta (Figure 7C-F). NID-1 puncta were distributed in a manner nearly indistinguishable from control animals in MEC-1[R1804C] mutants (Figure 7C-E, top row), while NID-1 puncta were denser in the MEC-1[C1808Y] animals (Figure 7C-E, bottom row, Figure 7F, Table S2).

**Figure 7.**
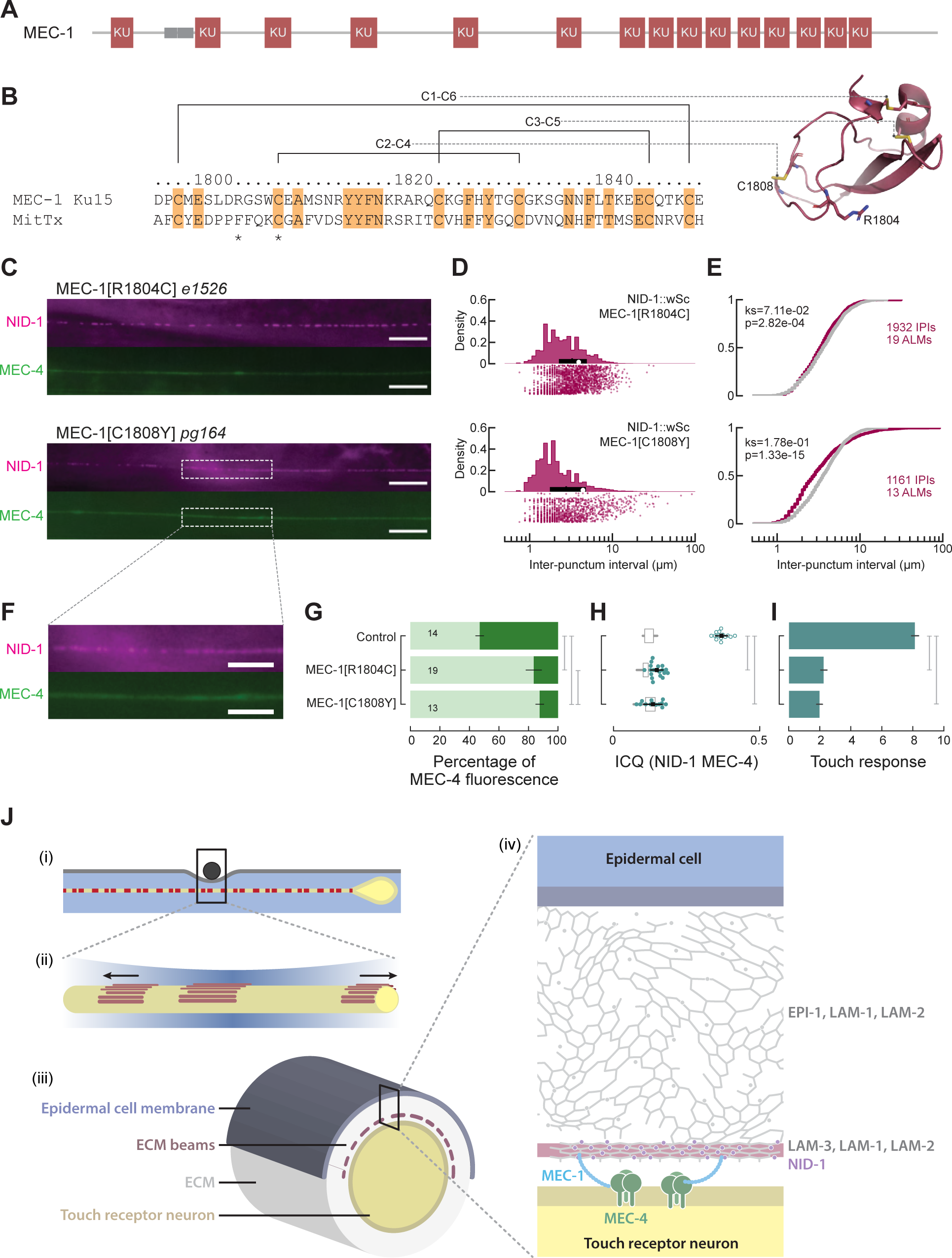
Disrupting Ku15 in MEC-1 causes misassembly of mechanosensory complexes and touch-insensitivity. **A.** Schematic diagram of the longest MEC-1 isoform showing fifteen predicted Kunitz domains (red) and EGF repeats (gray). **B.** Amino acid sequence of MEC-Ku15 compared to that of MitTx, a Kunitz domain-containing ion channel toxin (Baconguis, et al., 2014) (left) and a homology model (right) of MEC-1 Ku15 based on MitTx structures (PDB ID: 4NTX). Conserved amino acids are highlighted in orange. Asterisks denote the two mutated residues in MEC-1 Ku15, R1804 and C1808. **C.** Mechanosensory complexes visualized in dual-color transgenic animals expressing mNG::MEC-4 and NID-1::wSc in *mec-1* missense alleles affecting Ku15. Anterior is to the left, scale bar is 10µm. Boxed area in the MEC-1[C1808Y] image is enlarged in Panel F to show closely spaced NID-1::wSc puncta. **D.** Probability density distribution plots of IPIs (top) and the individual IPIs (bottom) for NID-1::wSc puncta in MEC-1[R1804C] (*e1526*) and MEC-1[C1808Y] (*pg164*) mutant animals. White circles denote mean, thin vertical lines denote the median and thick horizontal lines denote the interquartile ranges of the population. Summarized data are collected in Table S2. **E.** Cumulative distribution function (CDFs) of NID-1::wSc IPIs in the indicated *mec-1* alleles (magenta) vs control worms (gray). The results of two-sample Kolmogorov-Smirnov tests are indicated in each plot. See also Figure S7. **F.** Magnified view of the boxed area in C, showing NID-1 puncta and uniform MEC-4 fluorescence in a MEC-1[C1808Y] (*pg164*) mutant animal. **G.** MEC-4 distribution as a function of genotype for *mec-1* missense alleles. Stacked bars show the average relative fractions of mNG::MEC-4 fluorescence in the punctate (dark green) and diffuse (light green) pools. For each genotype, we analyzed 14, 19, and 13 independent replicates. One-way ANOVA shows significant differences in the MEC-4 diffuse fraction between genotypes [*F*(2,43) = 394.73, *p* = 2.15e-28]. Tukey-HSD post hoc analysis shows significant differences between all groups. Summarized data are collected in Table S2. **H.** Plot of ICQ values calculating MEC-4 and NID-1 colocalization as a function of genotype. Each circle (open circles: control data replicated from Figure 2F, filled circles: *mec-1* mutants) shows a measurement from a single ALM neurite. Black squares show the mean of the filled circles or open circles respectively, and the black bars show standard deviation. For every genotype except MEC-1[C1808Y], the ICQ of two fluorescence channels from the same cell (circles) are significantly higher (Student’s t-test, p<0.05) than that measured for simulated pairs (boxplots, showing the interquartile (box) and full ranges (whiskers), excluding outliers). One-way ANOVA shows significant differences in ICQ between genotypes [*F*(2,43) = 390.04, *p* = 2.74e-28]. Tukey-HSD post hoc analysis shows significant differences between control and both *mec-1* mutants but not between *mec-1(e1526)* and *mec-1(pg164)*. Summary data are in Table S2. **I.** Touch sensitivity as a function of genotype, assessed in a 10-touch assay. Bars represent the average (±SEM) of 75 animals tested in 3 independent cohorts blinded to genotype. One-way ANOVA shows significant differences between genotypes [*F*(2,222) = 280.68, *p* = 1.64e-61]. Tukey-HSD post hoc analysis shows significant differences between control and both *mec-1* mutants but not between *mec-1(e1526)* and *mec-1(pg164)*. Summary data are in Table S2. **J**. Proposed model of the relationship between MeT channels and ECM beam structures. (i) Schematic diagram of a TRN (yellow) embedded within the epidermis (blue), showing ECM puncta (red) as seen in fluorescence micrographs of NID-1, LAM-2 and LAM-2. Indentation is applied on the worm by a small bead (black) (ii) Expanded view of the indented region showing strain on the neurite (black arrows) and displacement of ECM beams (red). (iii) Schematic representation of a cross section of a TRN (yellow) and surrounding ECM (transparent gray) showing ECM beams (dark pink). (iv) Expanded view of the boxed area in (iii) showing spatial arrangement of the different proteins discussed in this manuscript. The EPI-1/LAM-1/LAM-2 laminin is shown as a loose hexagonal lattice filling most of the space between the epidermis (blue) and TRN (yellow). The LAM-3/LAM-1/LAM-2 laminin is shown as a tight and highly structured hexagonal lattice within the ECM beams (pink). NID-1 (purple circles) is also present in the ECM beam region. MEC-1 (cyan) connects the ECM beams to the MEC-4 channels (green) embedded in the TRN plasma membrane.

Laminin puncta were also retained in the MEC-1[R1804C] mutant animals and co-localized with nidogen (Figure S7). Collectively, these results demonstrate that an intact Ku15 is needed to link MEC-4 to BL structures. Additional support for this conclusion comes from the findings that, in these *mec-1* missense alleles, the bulk of mNG::MEC-4 fluorescence shifts to the diffuse pool (Figure 7G, Table S2), ICQ values for MEC-4/NID-1 are reduced (Figure 7H, Table S2), and animals are strongly touch-insensitive (Figure 7I). The findings from perturbations in Ku15 and their distinct effects on MEC-4, laminin, and nidogen puncta, are consistent with the hypothesis that MEC-1 physically binds to MEC-4 through Ku15.

## Discussion

Across animal taxa (Yin et al., 2021), the endings of vibrotactile sensory neurons are intimately associated with epidermal cells and basal laminae. Leveraging a suite of transgenic *C. elegans* expressing fluorescent-protein fusions from the endogenous loci of BL-encoding genes in the genome, we show that 12 conserved BL proteins are associated with TRNs in living animals. Among these, only laminin-β (LAM-1), laminin-γ (LAM-2) and nidogen (NID-1), display a punctate distribution along the TRNs but not in other tissues. Drawing upon still and time-lapse imaging of BL proteins and the MEC-4 ion channel subunit tagged alone and in pairs, FRAP studies, and genetic dissection, we provide evidence that laminin and nidogen co-localize with the MEC-4 channels responsible for touch sensation. These mechanosensory complexes are stationary and remarkably stable and we propose that they link the neuronal membrane to the extracellular matrix. At just over 1 µm long (Figure S2A) and separated by ∼4-5 µm (Table S2), on average, their dimensions are on par with the size and spacing of arrays of beam-like structures seen in ECM in serial-section TEM data stacks. These similarities prompt us to speculate that mechanosensory complexes associate with these structures (Figure 7J) and that the structures contribute to the immobility of the laminin, nidogen and MEC-4 puncta.

Consistent with this idea, MEC-4 redistributes to a less stable and more diffuse membrane pool in *mec-1* and *mec-9* mutants lacking discernable organized BL structures (Figure 3). Given that these mutants are completely touch insensitive (Du et al., 1996; Emtage et al., 2004), the MEC-4 proteins present in the diffuse membrane pool are not capable of supporting mechanosensation. A similar situation is evident in cultured TRNs physically separated from epidermal cells and their native BL structures. The few MEC-4 puncta that are present in these conditions seem to represent MEC-4 molecules that are sequestered in RAB-3-positive vesicles (Figure 3). Since cultured TRNs appear to maintain their cell-autonomous gene expression profiles (Zhang et al., 2002), their voltage-activated currents (Christensen and Strange, 2001), and their shape [this study and (Christensen and Strange, 2001; Christensen et al., 2002; Lockhead et al., 2016; Zhang et al., 2002), the failure to form mechanosensory puncta most likely arises from the physical separation from epidermal tissues and the subsequent disruption in the organization of laminin and nidogen along TRNs. The failure to maintain or recapitulate the *in vivo* organization of mechano-electrical transduction channels *in vitro* may have broad implications for studies of dissociated and cultured sensory neurons.

Loss of nidogen causes mild defects in behavioral and electrophysiological responses to touch (Figure 5H-J) and a substantial reduction in the density of MEC-4 and laminin puncta as well as the strength of their colocalization. This implies that the mechanosensory complexes retained in *nid-1* mutants (∼half of all MEC-4 puncta co-localize with LAM-1 and *vice versa*, Table S2, Figure S5) are fully functional and that nidogen plays a central role in their assembly or stability, but a minor role in their activation. It also implies that tactile sensing in *C. elegans* is robust, since nearly normal behavior is retained even though half of the MEC-4 channels are not properly aligned with their BL partners. By contrast, loss of MEC-7 ꞵ-tubulin causes severe defects behavioral and electrical responses to touch (Savage, et al., 1981; O’Hagan, et al., 2005), but modest decreases in the density of MEC-4 and nidogen puncta (Figure 4D-F) and their colocalization. Indeed, we found that more than two-thirds of MEC-4 puncta co-localized with NID-1 in *mec-7* mutants and *vice versa* (Table S2, Figure S4). Because MeT currents can be activated in *mec-7* mutants (O’Hagan, et al., 2005), however, the mechanosensory dysfunction in these mutants cannot be due to a complete loss of force transfer arising from disruptions to microtubules that also occurs in *mec-7* mutants (Chalfie and Sulston, 1981). Thus, more complex mechanisms or a combination of effects are needed to account for touch-insensitivity in this background, and any model would also need to consider the fact that *mec-7* decreases expression of the *mec-3* transcription factor and genes that require *mec-3* for their expression (Bounoutas et al., 2011). Thus, while *nid-1* mutants display essentially normal touch sensitivity with only half the channels aligned to BL puncta, *mec-7* mutants are effectively numb to touch despite retaining a significantly larger fraction of aligned channels. From these cases, we infer that proper organization of membrane-matrix complexes is necessary but that their proper placement is not sufficient by itself to support full vibrotactile sensation.

Laminin is a heterotrimeric molecule containing alpha, beta and gamma subunits (Aumailley, 2013), implying that one or both of the *C. elegans* alpha laminins also localize to the TRN-epidermal cell interface. Our experimental results are consistent with a model in which there are two distinct laminin networks bridging the gap between epidermal cells and the TRNs (Figure 7J). For instance, we found that EPI-1 uniformly coats the ALM (Figure 1) and that loss of *epi-1* function disrupts TRN attachment, but not the organization of MEC-4 puncta (Figure S5A-C). These findings suggest that an EPI-1/LAM-1/LAM-2 network coats the ALM uniformly at its interface with the ALM and is necessary for proper ALM ensheathment. They imply the presence of a uniformly distributed component of LAM-1 and LAM-2, in addition to the punctate component that we already described. This uniform component may be obscured in wild-type animals by the intense puncta and fluorescence from adjoining tissues. Indeed, LAM-2::mNG and NID-1::wSc were detected as a faint, but uniform fluorescence adhered to the ALM (Figure 3A, Figure S3B) in *mec-1(e1738)* and *mec-9(u437)* mutants lacking discrete laminin and nidogen puncta. Further, we posit that LAM-3/LAM-1/LAM-2 trimers form puncta along TRNs and that these laminin networks are necessary for MEC-4 puncta distribution. This model also predicts the presence of LAM-3 in a punctate distribution along the ALM, which we did not detect with fluorescently labeled LAM-3. New tools will be needed to test these predictions in detail and to discover other protein partners present in these networks. The proposed arrangement of two types of laminin networks would help to firmly link the ALM sensory ending to the surrounding epidermal cell.

We propose that sensory neurons instruct the assembly of mechanosensory complexes and discontinuous arrays of ECM beams seen in TEM data stacks. Two lines of evidence favor this idea. First, two genes expressed by the TRNs, *mec-1* and *mec-9,* are required to form aligned, stable complexes containing MEC-4, laminin, and nidogen. Both genes are predicted to encode secreted extracellular matrix proteins (Du et al., 1996; Emtage et al., 2004). Collectively, these observations suggest that these proteins could activate the assembly of discrete ECM structures. Second, ECM beams are formed adjacent to the AVM neuron in the ventral nerve cord at its interface with epidermal cells, but not neighboring neurons that share an interface with the same epidermal cell (Figure S1E). The MEC-1 protein plays additional roles. Prior work showed that it is required to attach the TRN to the epidermis and helps to ensure the proper positioning of its sensory neurite in the body (Emtage et al., 2004), but it is dispensable for force transfer from the skin to the TRN (Nekimken et al., 2020). Here, we establish that MEC-1 harbors specific domains required for co-assembly and alignment of MEC-4, laminin and nidogen. In particular, mutations predicted to disrupt folding of the C-terminal Kunitz domain in MEC-1 leave laminin and nidogen ECM structures intact but disrupt MEC-4 localization (Figure 7). Further experiments will test whether MEC-1 physically bridges the MeT channels with the ECM beams, as illustrated in Figure 7J, and whether these structures help transmit force to MeT channels.

Wherever neurons meet skin or muscle cells, the gap between them may well be occupied by BL proteins like laminin and nidogen. The BL is an active molecular medium, as evidenced by the fact that laminin plays an instructive role in organizing ion channels at the neuromuscular junction (Rogers and Nishimune, 2017) and instructing the formation of dendritic arbors (Yang and Chien, 2019). Laminin also governs the mechanosensitivity of cultured mouse dorsal root ganglion neurons (Chiang et al., 2011; Hu et al., 2010). In this study, we identified laminin and its binding partner, nidogen, as pivotal partners for organizing and immobilizing the mechano-electrical transduction channel, MEC-4, along *C. elegans* TRNs. Two other sensory mechano-electrical transduction channels also localize to discrete subcellular positions in fixed samples: NOMPC (Lee et al., 2010; Liang et al., 2011) and Piezo2 (Handler et al., 2023). These channels also appear to localize at the interface between sensory nerve endings and skin cells, although their stability, dynamics, and positioning relative to the BL is not yet known. Our findings extend the role of the BL from structure to function and defines a mechanosensory apparatus as a multiprotein complex linking the neuronal membrane to the BL matrix.

## Limitations of the study

This study characterizes electron-dense structures and mechanosensory protein complexes at the sensory neuron-epidermal cell interface. The complexes are essential for touch sensation and contain both neuronal MeT channels and extracellular assemblies of laminin and nidogen; they depend on secreted neuronal factors proposed to transform continuous basal lamina into discrete, stable, and stationary complexes. However, there are limitations that require further investigation or new tools. Despite conducting a comprehensive visual screen of more than two dozen ECM proteins, it is likely that additional proteins are present at the sensory neuron-epidermal cell interface. Of the proteins we analyzed, one tagged protein was not visible in any tissue in adult animals (LAM-3 α-laminin) and another tagged protein caused defects in touch sensation (FBL-1). Tagging MEC-1 and MEC-5 at their endogenous loci disrupted touch sensation, limiting our ability to re-examine the location of these proteins and to investigate their interaction with *mec-4* and *nid-1* mutants. The widespread distribution of the visible laminin subunits (LAM-1, LAM-2, EPI-1) contributed to background fluorescence that affected our analyses of their distribution and colocalization with the MEC-4 ion channel. Additionally, we could not systematically study how loss of laminin affects mechanosensory complex assembly, stability, or function since loss-of-function mutations in all four laminin genes cause embryonic or larval lethality. Biochemical studies and purification of this ion channel-ECM complex from *C. elegans* are hampered by the scarcity of channel proteins relative to ECM proteins. Identifying the protein partners that comprise nanostructures such as the ECM beams visible in serial-section TEM structures is notoriously difficult and may require new technical approaches. The anatomy and molecular physiology of the *C. elegans* TRNs differs from other somatosensory neurons with respect to their position within the skin and the ion channels responsible for mechanotransduction. Therefore, it may be challenging to immediately apply these results to other sensory neurons and organisms, including those already known to share an intimate association with epidermal cells. Despite these limitations, this study is a first step toward deciphering the active nature of the interaction between sensory neurons and skin.

## Acknowledgements

We thank Tom Clandinin, Ellen Kuhl, and Lucy Wang for comments and advice; the entire WormsenseLab for feedback on experimental and analytical design; Zhiwen Liao for technical support and worm injections; Juan Cueva for pilot studies, quantification of ECM beams in ssTEM datasets, and 3-D visualization of ECM beams; and Richard Fetter for contributing independently generated ssTEM datasets. Martin Chalfie, Kang Shen, David Sherwood, and the Caenorhabditis Genetics Center (supported by the NIH Office of Research Infrastructure, P40 OD010440) provided *C. elegans* strains. We are especially grateful to David Sherwood for sharing key transgenic strains prior to publication. Confocal fluorescence imaging and legacy ssTEM datasets used instruments in the Stanford University Cell Sciences Imaging Core Facility (RRID:SCR_017787) and protein micropatterning used instruments in the Stanford Nanofabrication Facility (supported by NSF Award ECCS-2026822). Additional ssTEM images were acquired at the Nanoscale Biomedical Imaging Facility, Toronto. Research funded by an NIH grant R35NS105092 to MBG, CIHR foundation scheme 154274 to MZ, F99NS115219 fellowship to JAF, and a F32NS116193 fellowship to DC.

## Author contributions

Conceptualization, A.D., J.F., B.M., D.C., M.B.G.;

Methodology, A.D., J.F., B.M., L.W., C.J.; Software, A.D., J.F., L.W.;

Validation, A.D., J.F., L.W.;

Formal analysis, A.D., J.F., B.M., L.W.;

Investigation, A.D., J.F., B.M., L.W., D.C., C.J.;

Resources, A.D., J.F., L.W., B.P., E.K., M.Z., M.B.G.;

Data Curation, A.D., J.F., B.M., L.W.;

Writing - Original draft, A.D., J.F., L.W., D.C.;

Writing - Review and editing, A.D., J.F., L.W., D.C., B.M, M.Z., M.B.G.;

Visualization, A.D., J.F., L.W.;

Supervision, A.D., M.Z., M.B.G.;

Project administration, A.D., M.Z., M.B.G.;

Funding acquisition, M.Z., M.B.G.

## STAR Methods

### RESOURCE AVAILABILITY

#### Lead contact

Further information and requests for resources and reagents should be directed to and will be fulfilled by the lead contact, Miriam B. Goodman.

#### Material availability

*C. elegans* strains newly generated for this paper are available from the lead contact upon request. Interested parties should submit requests to the lead contact indicating the strain and genotype of each of the required *C. elegans* strains.

#### Data and code availability

This paper did not generate new data conforming to standardized data types, including nucleotide sequences, proteomics, metabolomics, or structures of biological macromolecules. This paper did generate new *C. elegans* genotype-phenotype relationships, fluorescence images, electrophysiological recordings, and behavior testing results.

● Genotypic and phenotypic information about new *C. elegans* alleles created in this study will be deposited to www.wormbase.org.
● All original code has been deposited at GitHub and is publicly available as of the date of publication. DOIs are listed in the key resources table.

### EXPERIMENTAL MODEL AND SUBJECT DETAILS

#### C. elegans

This study uses wild-type, mutant and transgenic *C. elegans* nematodes. Only hermaphrodites were analyzed in this study, although males were used to perform genetic crosses. A complete list of all *C. elegans* strains used in this study with their genotypes and provenance is available in the Key Resources Table. The list includes transgenic and mutant lines generated specifically for this study, obtained from prior work, or from the Caenorhabditis Genetic Center. Animals were maintained at 20°C on NGM agar plates seeded with *E. coli* OP50, unless otherwise mentioned.

All animals studied by light microscopy, tested in behavioral assays, or subjected to *in vivo* electrophysiology were age-synchronized adopting standard methods (Stiernagle, 2006). In brief, we harvested gravid adults in ∼1 mL of water and obtained eggs from these animals by treating them with household bleach (100µL) and KOH (20µL, 5M solution). Eggs were transferred to NGM agar plates seeded with bacterial food and allowed to grow into young adults (∼3 days at 20°C).

#### Primary cells

**(nematodes)** As detailed below, we generated cultures of primary cells from transgenic *C. elegans* nematodes expressing fluorescent proteins. Cells were plated on coverslips coated uniformly by peanut lectin or bearing thin stripes of peanut lectin fabricated in-house as described below (protein patterning coverslips) and maintained in a benchtop chamber at room atmosphere and temperature (22-24°C) for 1-7 days.

### METHOD DETAILS

#### Gene editing via CRISPR/Cas9 techniques

##### Tagging endogenous proteins

Unless indicated otherwise, we relied on homologous recombination via the self-excising cassette (SEC) method (Dickinson et al., 2015) to insert fluorescent protein-encoding DNA at four endogenous loci: *lam-1, lam-3, epi-1* and *nid-1*. We created a template repair plasmid (mg682) with wormScarlet (wSc) based on the SEC repair plasmid designs described in Dickinson et al., 2015. We cloned in left and right homology arms corresponding to the insertion sites at the *lam-1, lam-3, epi-1* or *nid-1* loci into this template plasmid by Gibson assembly.

Repair plasmids and small-guide RNA (sgRNA) targets were designed based on data curated by WormBase (Davis et al., 2022). To generate each strain, we injected a plasmid mixture into the gonads of 40-60 young adult animals. The mixture contained plasmids encoding the following elements: Cas9-sgRNA (50 ng/µL), repair template (50 ng/µL), visible markers [pCFJ104 (myo-3p::mCherry), 5 ng/µL; pCFJ90 (myo-2p::mCherry), 2.5 ng/µL]. Each injected animal was allowed to lay eggs for 2-3 days at 25°C, in the absence of selection. To select for progeny carrying edits, we added hygromycin (500 µL, 5 mg/mL) to each plate and allowed animals to grow at 25°C for an additional 4-5 days. Candidate transgenic animals survived hygromycin treatment, lacked red fluorescent extrachromosomal array markers, and displayed a Rol phenotype conferred by the insertion of the SEC carrying the *sqt-1(e1350)* mutation (Dickinson et al., 2015). To remove the SEC, we heat-shocked plates containing ∼6 L4 rollers (37°C, 75 minutes) and allowed the resulting progeny to grow at 20°C for 3-4 days. Single, adult NonRol animals (worms that lost both copies of the SEC) were transferred to growth plates and allowed to generate progeny. These candidate transgenic animals were evaluated by visualizing fluorescence, PCR genotyping, and sequencing. Homozygous wSc-tagged *lam-1* animals were sterile and exhibited several morphological defects, similar to those previously reported for a mNeonGreen(mNG)-tagged *lam-1* (Keeley et al., 2020). Consequently, we maintained and imaged these animals as viable heterozygotes.

We used a slightly modified approach from above for creating the PAT-2::wSc strain. In this approach we used a template repair plasmid in which the Lox2272-site flanked selection cassette contained the Hygromycin resistance gene, but lacked the *sqt-1(e1350)* reporter and the heat-shock inducible Cre genes (mg546). The repair plasmid was designed such that presence of the selection cassette will prevent translational fusion of the wSc tag at the PAT-2 C-terminus. After confirming insertion of the wSc tag and the selection cassette at the *pat-2* locus by PCR, we injected the worms with a plasmid encoding *eft-3* promoter driven Cre, alongwith a co-injection marker (pCFJ90 (myo-2p::mCherry), 2.5 ng/µL). We then screened for the loss of the selection cassette among the progeny of the injected worms, by looking for expression of wSc tagged PAT-2 in all body wall muscle cells as well as a loss of the *myo-2p*::mCherry red pharynx marker.

##### Mutating endogenous gene loci

To generate *C. elegans* strains carrying genetic deletions or point mutations in *nid-1*, *lam-2* or *mec-1* loci, we applied the co-CRISPR method (Arribere et al., 2014; Paix et al., 2015), using *dpy-10(cn64)* as the co-CRISPR target. This work relied on genetic and genomic data curated and maintained by WormBase (Davis et al., 2022). All reagents for this purpose were purchased from IDT Technologies: Cas9 protein (Alt-R® S.p. Cas9 Nuclease V3, 100 µg, Catalog # 1081058), tracrRNA (Alt-R® CRISPR-Cas9 tracrRNA, 5 nmol, Catalog # 1072532), crRNA for each loci (Alt-R® CRISPR-Cas9 crRNA, 2 nmol), single stranded oligomeric DNA repair templates (Ultramer™ DNA Oligonucleotides). We prepared injection mixes with equimolar amounts of tracrRNA, crRNA and Cas9 nuclease, such that the final concentration of each was 1.5 µM. If multiple crRNAs were used for a single target site, the amounts were adjusted such that the total crRNA concentration was 1.5 µM. The *dpy-10* crRNA was used at a ratio 1:14 to the target crRNAs. In the first step, we allowed the tracrRNA and crRNAs to anneal at 95°C for 5 min, followed by incubation at room temperature for 5 min. Next, we added the annealed RNA mixture to Cas9 and incubated this solution at 37 °C for 10 min. Finally, ssDNA repair template was added to generate a final concentration of 0.5 µM and the total volume was adjusted to 20 µL using sterile water. For each gene editing round, we injected 15-20 young adult animals and transferred individual animals to growth plates for 3-4 days at room temperature. Plates that contained many Rol progeny as evidence of successful editing were selected for further analysis. Specifically, we isolated single Rol animals and allowed them to lay eggs for 2 days (room temperature), after which the roller animals were screened for the desired mutation using PCR. Non-roller progeny animals from the positive plates were then isolated and screened in the same way. The target loci were sequenced in candidate lines to confirm the presence of the desired mutation and validate the gene-edited line.

##### Primary nematode cell culture

We generated cultures of mixed embryonic *C. elegans* cells following previous methods (Bianchi and Driscoll, 2006; Sangaletti and Bianchi, 2013; Strange et al., 2007) and introduced modifications that improved yield. In brief, the procedure involves treating suspensions of isolated embryos with chitinase to soften the egg shell, mechanical trituration and filtration to dissociate cells and remove debris, followed by plating onto treated glass coverslips and culturing at room temperature and atmosphere for 1-7 days.

To improve yield, we first sought to improve age-synchronization and to increase the number of embryos available for cell isolation. In particular, we grew animals in three cycles on 10-cm plates. During the first cycle, we used standard hypochlorite bleaching methods (6% bleach, 150 mM NaOH: Stiernagle, 2006) to harvest embryos from animals grown on standard NGM plates seeded with OP50 *E. coli.* These embryos were transferred to 10-cm enriched peptone plates seeded with NA22 *E. coli* and allowed to grow into gravid adults. This process was repeated for two additional cycles. Following these steps, we harvested embryos after separating gravid adults from bacteria through several centrifugation and washing cycles and sodium hypochlorite treatment to harvest embryos. The resulting embryos were rinsed three times with 1x egg buffer (25 mM HEPES, 118 mM NaCl, 48 mM KCl, 2 mM CaCl2, 2 mM MgCl2, adjusted to a pH of 7.3 with NaOH). For quality control purposes, we measured the osmolality of the egg buffer with a freezing-point depression osmometer (AdvancedInstruments) and used batches that had an osmolality of 340 ± 5 mOsm.

Next, we separated eggs from lytic fragments of older worms by resuspending the pellet in a sucrose solution (60% w/v sucrose in deionized water) and centrifugation at 350g for 5 minutes. Following these steps, we incubated embryos in chitinase (25 U/mL) while rocking until ∼80% of eggshells had been digested (∼50 min), an endpoint that we confirmed visually. Finally, we rinsed dissociated embryos and dissociated them mechanically by trituration first through a sterile 21-gauge needle and then through a sterile 25-gauge needle. We used a 5-µm filter to remove debris from the resulting cell suspension, rinsed and resuspended the cell suspension in sterile L-15 cell culture media without HEPES or phenol red (Gibco^TM^, Catalog# 21083027), and seeded the cells onto glass coverslips either coated uniformly with peanut lectin or bearing micropatterns of peanut lectin. These procedures were performed on a standard benchtop using aseptic technique.

##### Protein micropatterning glass coverslips

To generate thin stripes of adhesive proteins, which constrained the shape of *C. elegans* neurons in culture, we used a light-activated molecular adsorption technique and a commercial instrument (Primo, Alvéole USA, Fairfax, VA, USA) to generate protein micropatterns on glass coverslips. We used open-source graphic design software (Inkscape) to design custom patterns consisting of thin (8-pixels or 2.2 µm) stripes on 1824×1140 pixel canvas size. Stripe width and spacing was optimized to enable touch receptor neurons to elaborate straight neurites that were confined to lateral strips and did not traverse areas lacking peanut lectin (PNA), as described (Franco, 2021). We used both unlabelled peanut lectin and Alexa Fluor™ 594-conjugated lectin PNA.

We adapted a protein micropatterning approach (Strale et al., 2016), as follows. For convenience and simplicity, we used glass-bottom six-well tissue culture-grade microtiter plates (6 Micro-well glass bottom plate with 20 mm micro-well #1.5 cover glass, Cellvis ™, Catalog# P06-20-1.5-N) for cell culture experiments involving patterned coverslips. First, we plasma-treated the plates with ionized atmospheric gas for 1 minute (25 forward, zero reflected power, Plasmatech P-50). Next, we positioned a PDMS reagent basin in each well, added poly-l-lysine or PLL (0.01% in sterile PBS, ∼35 µl) to each well, and allowed the PLL solution to incubate for 1 hour. Following this step, we added mPEG-SVA (100 µg/mL, Laysan Bio Inc.) freshly prepared in pH-adjusted 100 mM HEPES, pH 8.0 buffer and incubated the coverslips for an additional 1 hour. We fabricated reagent basins from PDMS sheets in-house using a consumer-grade, computerized die-cutting machine (Silhouette, Cameo, Lindon, Utah, USA). Lastly, we rinsed the coverslips with distilled water and allowed them to air dry prior to the application of proprietary photoactivatable reagent, PLPP gel (Alvéole, diluted in pure ethanol at 1:6 volumetric ratio).

We allowed coverslips to air dry prior to patterning with the Primo instrument (25 ms exposure, 100% laser power, 50 mJ/mm^2^ dosage), which uses light to degrade the PLPP gel according to the user-defined pattern. We transferred peanut lectin protein (PNA) to patterned coverslips in two incubation steps: 1) PBS for >5 minutes; 2) PNA (500 µg/mL) for 20 minutes. which enabled us to visualize the patterns. We fabricated patterns one day prior to plating dissociated embryonic *C. elegans* cells and kept them hydrated with sterile PBS until use.

##### *C. elegans* touch assay

We measured touch responses using a ten-touch assay (Chalfie et al., 2014; Vasquez et al., 2014). Specifically, we delivered mechanical stimuli to age-synchronized, well-fed young adult animals with an eyebrow hair glued to a toothpick touching animals alternately in anterior and posterior regions of the body for a total of 10 times. Animals were scored as responding if they reversed direction following each stimulation. This method delivers supra-threshold stimuli, independent of experimenter or the specific eye-brow hair used (Nekimken et al., 2017). We analyzed touch sensitivity of groups of 4-5 strains and used wild-type (N2) and TU253 *mec-4(u253)* null mutants as touch-sensitive and touch-insensitive (Mec) references, respectively.

Animals were tested blind to genotype in cohorts of 25 animals. At least three independent cohorts were tested for each genotype or condition. For strains containing *lam-1(pg136)* allele (LAM-1::wSc), we picked ca. 50 heterozygous animals (identified by dimmer red fluorescence, normal locomotion and body morphology compared to homozygous *lam-1(pg136)* animals which were sluggish, dumpy and/or had severely malformed internal organs) to fresh plates a few hours prior to touch testing.

##### Whole-cell patch clamp electrophysiology

We recorded from the ALM neurons in TU2769 *uIs31 [mec-17p::GFP]* and GN932 *uIs31; nid-1(cg119)* lines, following established procedures (Eastwood et al., 2015; Katta et al., 2019; O’Hagan et al., 2005). As a consequence of geometric constraints imposed by our stimulator system, most of our recordings are from the ALMR neuron. The ALMR and ALML neurons are bilaterally symmetric and exhibit no detectable differences in voltage- or touch-evoked currents (this study and Eastwood et al., 2015; Katta et al., 2019; O’Hagan et al., 2005). We used the following salines in all recordings. Extracellular saline contained (in mM): NaCl (145), KCl (5), MgCl2 (5), CaCl2 (1), and Na-HEPES (10), adjusted to a pH of 7.2 with NaOH. We added D-glucose (final concentration of 20mM) to external saline, bringing the osmolarity to ∼325 mOsm. Intracellular recording solution contained (in mM): K-gluconate (125), KCl (18), NaCl (4), MgCl2 (1), CaCl2 (0.6), K-HEPES (10), and K2EGTA (10), and adjusted to a pH of 7.2 with KOH. We added a fluorescent dye, sulforhodamine 101 (1mM, Invitrogen), to the intracellular solution in order visually confirm whether or not the whole-cell configuration was achieved in each recording. We used an EPC-10 USB amplifier controlled by Patchmaster software (version 2×90; HEKA/Harvard Biosciences) to record membrane current, deliver voltage-clamp stimuli, and to control the mechanical stimulator. Mechanical stimuli were delivered using an open-loop system adapted from (Peng and Ricci, 2016) and described previously (Katta et al, 2019). Analog data (membrane potential, membrane current) were filtered at 2.9 kHz and digitized at 5 kHz.

Voltage-activated currents were measured in response to 100ms pulses between ™80 and 80mV (in 20mV increments) from a holding potential of ™60mV. We applied each voltage protocol in triplicate and averaged the resulting membrane current traces to improve the signal to noise, as described (Goodman et al., 1998). Membrane voltage was corrected for errors due to liquid-junction potentials (−14 mV) and residual series resistance measured in each recording. We omitted recordings with series resistance larger than 200 MΩ and holding current more than ™10 pA at ™60 mV. MRCs were evoked by indentation steps (300 ms) with an interstimulus interval of 1s from a holding potential of ™60mV, as described by Katta et al., 2019. We applied 3 repetitions of mechanical stimuli and averaged the resulting MRC traces. We used peak-finding software to measure peak MRC currents at the onset and offset of the mechanical stimulation pulse. For clarity, only the ‘on’ MRCs are shown in this study.

##### Protein structure homology modeling

We generated a homology models for the NID-1[G3]-LAM-2[EGF8] complex and MEC-1[Ku15] using the Phyre2 protein fold recognition server (Protein Homology/analogY Recognition Engine V 2.0) (Mezulis et al., 2015). For the NID-1[G3]-LAM-2[EGF8] complex model, we used the crystal structure of mouse nidogen bound to laminin-γ (PDB ID: 1NPE; Takagi et al., 2003). For the MEC-1[Ku15] model, we used the structure of the neurotoxin MitTx-alpha bound to the chicken ASIC1 ion channel (PDB ID: 4NTX; Baconguis et al., 2014). Both models were rendered for publication using the The PyMOL Molecular Graphics System, Version 2.5.1 Schrödinger, LLC.

##### Transmission electron microscopy

We relied on multiple, independent serial-section TEM (ssTEM) datasets to investigate the ultrastructure of ECM structures associated with the TRNs in wild-type animals. In all cases, animals were prepared for TEM by high-pressure freezing followed by freeze substitution with chemical fixatives and embedded in plastic resin. The image series of ALM and AVM neurons shown in Figures 1E and Figure S1A-B and E(left) is derived from a wild-type (N2), L3 stage larva, while those of PLM and PVM neurons shown in Figures S1C-D and E(right) is from a wild-type (N2), L4 stage larva. We estimated ECM beam dimensions and distance from the TRN plasma membrane and basal membrane of epidermal cells from ssTEM images of ALM neurons prepared in young adult worms, prepared for a prior publication (Kreig et al., 2017). Despite variations in sample preparation, life stage, embedding resins, sample preparation and microscope instrumentation, ECM beam structures have similar appearances and structures in all ssTEM datasets we have examined.

For L3 larval worms, we relied on high pressure freezing followed by freeze substitution, as previously described (Mulcahy et al., 2018; Witvliet et al., 2021). Briefly, mixed-stage wild-type (N2) animals were collected into 3 mm high pressure freezing carriers with 100 μm wells (Leica, product #16770141), using OP50 *E. coli* as a filler. A flat lid (Leica, product #16770142) was placed on top and the sandwich was high pressure frozen using a Leica EM ICE (product #16771801). Freeze substitution was accomplished by ramping the sample from ™90 to ™60°C over 20 hours in 0.1% tannic acid and 0.5% glutaraldehyde, followed by washing 4 times with acetone over 3 hours, exchanging with 2% OsO4 and ramping the temperature to ™20°C over 8 hours, holding at ™20°C for 11 hours, ramping the temperature to 0°C over 4 hours and washing 4 times with acetone over 1 hour. We then infiltrated the sample with Spurr-Quetol resin, embedding single animals, and cured the samples (60°C, 24 hours). The age of this sample was determined by the lineage of neuronal and non-neuronal cells in the volume (Witvliet et al., 2021). We collected 50 nm serial sections on 2×0.5 mm slot grids using a Leica UC7 ultramicrotome and 35° diamond knife (Diatome). Sections were post-stained with 2% aqueous uranyl acetate and 0.1% Reynold’s lead citrate, and imaged at 0.768 nm/pixel X-Y resolution using a Technai T20 TEM at 120kV equipped with a digital camera (Gatan Orius). We aligned and down-sampled 212 and 427 serial TEM micrographs to 3.072 nm/pixel resolution, estimated to cover 10.6 μm (ALML), 10.6 μm (ALMR), and 21.53 μm (AVM) processes.

For L4 larval worms, samples were high-pressure frozen in M9 media containing 20% BSA and E. coli in specimen carriers with a 50 µm deep well (Cat. no. 390, TechnoTrade International, Inc.). These specimen carriers were covered with the flat side of a Type B specimen carrier (Cat. no. 242, Technotrade International Inc.) and frozen with a BalTec HPM01 high pressure freezer (Liechtenstein). Freeze-substitution was carried out with a Leica AFS2 unit in acetone containing 1% OsO4, 0.1% uranyl acetate, 1% methanol, and 3% water (Walther and Ziegler, 2002). Samples were held at ™90°C for 16 hours, ramped to ™60°C in 6 hours, held at - 60°C for 6 hours, ramped to ™25°C in 7 hrs, held at ™25°C for 6 hours, and finally ramped to 0°C over 5 hours. At the completion of substitution, we rinsed samples in acetone, infiltrated and then polymerized in Eponate 12 resin (Ted Pella, Inc.). We cut serial 50 nm cross-sections with a Leica UCT ultramicrotome using a Diatome diamond knife, picked up the sections on Pioloform-coated slot grids and stained with uranyl acetate and Sato’s lead (Sato, 1968). We imaged the sections with an FEI Tecnai T12 TEM at 120 kV equipped with a Gatan U895 4k x 4k camera (Gatan, Pleasonton, CA) at a pixel resolution of 1.01 nm/pixel using SerialEM (Mastronarde, 2005).

#### *Light* microscopy

##### *C. elegans* animal mounting for imaging

To visualize and analyze MEC-4 and protein constitutions of the basal lamina in living animals, we generated sets of still and time-lapse images. We imaged *C. elegans* animals mounted on agarose pads (5% wt/vol) on glass slides. In brief, we used levamisole (5mM in M9 buffer) to immobilize age-synchronized young adult animals or L1 larvae. We mounted young adult animals on grooved agar pads fabricated by casting the pads on the surface of a standard LP record (Rivera Gomez and Schvarzstein, 2018). This approach generates worm-sized grooves and constrains animals to a straight posture within the grooves, enabling us to image dozens of animals in a single imaging session. This approach was not appropriate for L1 larvae, however, due to their narrower bodies. Instead, we collected synchronized L1s in M9 buffer, treated them with levamisole for ∼10 minutes until most animals assumed a straight posture, concentrated the animals by centrifugation, and transferred 2 µL of concentrated L1s to surface of a smooth agarose pad bearing a small droplet of levamisole (5mM in M9). To keep the samples moist during each imaging session, we covered all mounted animals with a glass cover slip and sealed the coverslip with VALAP (1:1:1 <u>V</u>aseline:<u>La</u>nolin:<u>P</u>araffin).

##### Wide-field and confocal fluorescence microscopy (still images)

We generated wide-field fluorescence images of the ALM neurites and associated basal lamina (BL) structures using an automated inverted compound microscope equipped for epifluorescence imaging (Keyence BZ-X800). Using the instrument’s stitch and align function, we first collected a low magnification image (4X) and selected individual animals to image at high magnification (60X, Nikon Plan Apo lambda, oil immersion, NA=1.40). In most cases, we used the stitch and align function to generate complete images of the ALM neurite. For all experiments (except those using strains expressing mCherry::RAB-3) designed to compare puncta distribution or fluorescence intensity across genotypes, we kept the following image acquisition parameters constant: exposure time (mNG::MEC-4 = 1 s, LAM-1::wSc = 0.2 s, LAM-2::mNG = 1.2 s, NID-1::wSc = 0.133 s), binning (1×1), camera gain settings (+6dB). We acquired images of strains expressing the synaptic vesicle marker, mCherry::RAB-3 using 2×2 binning, and the following exposure times: mNG::MEC-4 = 0.2 s, mCherry::RAB-3 = 0.1 s. For imaging the movement of MEC-4 in vesicles over time (Figure S3C,S3E), we used time-lapse software to collect dual-color time-lapse image stacks using Keyence BZ-X800 and Nikon Plan Apo lambda 60x objective, using the following acquisition parameters: 3×3 binning, 1 frame every 3s, for 40 frames (or a total duration of 120s or 2 minutes). To improve visualization of TRN-associated BL structures, we generated confocal stacks of transgenic animals (Figure 1C- D) using an inverted, laser-scanning confocal microscope (Leica SP8, Stanford Cell Sciences Imaging Facility) and a 40X oil-immersion lens (Leica 40X HC PL APO, CS2, oil, NA=1.30).

##### Time-lapse imaging and fluorescence recovery after photobleaching

**(FRAP)** We conducted FRAP experiments in living animals, mounting animals as described above and generated time-lapse images on an inverted, laser-scanning confocal microscope (Leica SP8, Stanford Cell Sciences Imaging Facility) at 40X magnification (Leica 40X HC PL APO, CS2, oil, NA=1.30). Briefly, an image scientist (AD or DC) acquired images at 1 fps for 10 frames prior to bleaching, selected a region of interest (∼15×15 pixels) containing a defined punctum or in an inter-punctum interval, and scanned that region at maximum power for 3.5 s to bleach the fluorophores present. Following bleaching, we acquired images every 5 seconds (mNG::MEC-4, LAM-2::mNG, NID-1::mNG and SDN-1::mNG) or every 1 second (myr::GFP) for 60 frames.

##### C. elegans TRNs in culture

We used the Keyence instrument (BZ-X800) to generate still fluorescence images of dissociated, live TRNs using a similar strategy, 24 or 48 hours after plating on 6-well glass-bottom dishes coated with PNA, uniformly or in a striped pattern. We acquired images using a 60x oil immersion objective (Nikon Plan Apo lambda 60x), using the following parameters: binning (1×1), 1.5 ms exposures in the green (mNG) channel and 16ms exposures in the red (mCherry) channel. In addition to epifluorescence images, we also collected brightfield images of the same fields (1×1 binning, 6 msec exposure). Widefield images were saved as 32-bit RGB*.tiff files. We used time-lapse software to collect dual-color time-lapse image stacks (Figure S3D) using the same microscope (Keyence BZ-X800) and objective (Nikon Plan Apo lambda 60x), using the following acquisition parameters: 1×1 binning, 1 frame every 6s, for 20 frames (or a total duration of 120s or 2 minutes).

### QUANTIFICATION AND STATISTICAL ANALYSIS

#### Serial-section TEM reconstruction

For data stacks generated for Krieg, et al., 2017, we used ImageJ (Rasband, 1997) to measure the width and height of ECM beam structures in single transmission electron micrographs. In order to determine their length and generate three-dimensional models, we used Reconstruct software (Fiala, 2005), a legacy editor for serial-section TEM analysis running under WindowsXP or Linux (https://www.bu.edu/neural/Reconstruct.html). In brief, all images of consecutive sections were aligned manually and the outline of the ECM beams, TRN membrane, and epidermal cell basal membrane were traced by hand. For all other datastacks, either individual images or montaged sections were aligned with TrakEM2 (Cardona et al., 2012) in Fiji (Schindelin et al., 2012) using elastic deformation corrections and examined them visually. The length of ECM beam arrays and the distance separating adjacent arrays were determined by observing the presence of beams in individual sections in these datastacks.

#### Fluorescence signal analysis in ALM neurons

##### Pre-processing fluorescence images of neurites

We used Fiji (Schindelin et al., 2012) and Python to build an efficient, reproducible image processing pipeline for analyzing fluorescence signal distribution along ALM neurons. As described above, we first generated images showing the entire ALM neurons using the stitch- and-align function of the Keyence microscope proprietary software and exported images in *.tif format. Next, we traced the anterior processes of the ALM neurites from the cell body to the distal tip (unless otherwise mentioned) as a segmented line (line parameters: width = 20 pixels, spline fit). We converted these traces to straightened cropped sections using the built-in ‘Straighten’ function in FIJI.

##### Puncta analysis

We then analyzed the straightened neurite images generated as described above by a custom Python script (https://github.com/wormsenseLab/Neurite_fluorescence_analysis), handling background subtraction and calculating the fluorescence intensity of a labeled protein along the neurite. Puncta were detected from these plots using the ‘find_peaks’ function from scipy library. To minimize bias in peak detection from differences in expression levels, we employed adaptive threshold estimation for each image, as a function of the local minima and global mean and standard deviations of the fluorescence intensity for each image. For MEC-4 images, the threshold estimation function additionally took into account the local average intensity since there was often a gradual decrease in overall intensity from the proximal to the distal end of the neurite. We recorded the position of each puncta as the distance of the puncta peak from the cell body along the neurite and calculated inter-punctum intervals as the distance between the peaks of neighboring puncta. We also calculated the punctate and diffuse fluorescence fractions from these intensity profiles. Except for analysis of mCherry::RAB-3, we analyzed fluorescence in the entire ALM neurite. For strains expressing mCherry::RAB-3, we focused analysis on a 200 µm-long segment of the ALM neurite immediately anterior to the cell body.

##### Colocalization analysis

We analyzed colocalization of two proteins along the ALM neurite by calculating the Intensity Correlation Quotient (ICQ). ICQ is a measure of pixel-by-pixel co-variation of intensities between two fluorescent channels. It is defined as the ratio of the number of pixels where the product of difference from mean for the red and green channels are positive, to the total number of pixels subtracted by 0.5 (Li et al., 2004). As a consequence, the ICQ values vary from 0.5 (perfect co-localization) to ™0.5 (perfect exclusion). Dual-color labeling that is uncorrelated or whose analysis is impeded by background noise will give a value close to zero. To validate this approach to colocalization in the ALM neurons, we generated reference conditions in which near-perfect co-localization was expected (*i.e.* labeling NID-1 with mNG and wSc) and in which no colocalization was expected (*i.e.* dual-labeling with a punctate and diffuse marker). Previously, we used this approach to analyze colocalization of antibody-labeled MEC-4, MEC-2 and MEC-5 in fixed TRNs, using two antibodies directed against MEC-2 as a co-localization reference (Cueva et al., 2007). As an additional means of cross-validation, we randomly selected red and green channels from different cells within each dataset and used the resulting simulated pairs to compute ICQ. In all cases and as expected, the simulated pairs had ICQ values close to zero (Figure 2F, 7H, S4E, S5E, S7G, S7J).

We also determined object colocalization by measuring the percentfage of detected puncta in one channel that co-localized with puncta detected in the other channel and vice-versa. We defined positive co-localization if the peaks of two puncta were less than 0.5 um apart. As above, we also computed the co-localization of puncta between red and green channels randomly selected from different cells within each dataset, to estimate the expected baseline level of puncta colocalization between two random fluorescence profiles with similar overall number and spacing of puncta. In all cases and as expected, these values were close to zero (Figure S2F-G, S4F-G, S5F-G, S7H-I, S7K-L)

##### FRAP analysis

We processed the time-lapse image stacks of ALM neurons before and after photobleaching as follows. First, we traced ALM neurites (using the segmented line tool in ImageJ), converting the image stack to a kymograph of the ALM neurite. We then inspected kymographs for global movement of the neurite during image acquisition and for the appearance of moving puncta, excluding samples with clear evidence of movement from further analysis. Once validated, we generated a straightened, 21-pixel wide cropped image stack, containing the ALM neurite. We then analyzed the cropped image stacks using a custom Python script (https://github.com/wormsenseLab/FRAP) to detect the bleached region based on the expectation that it would contain the maximum change in fluorescence intensity between the last three frames collected prior to bleaching and the first three frames after bleaching. The average fluorescence intensity in the bleached region for each time point was corrected for overall photobleaching over the time period of image acquisition, by dividing by the average fluorescence intensity in the non-bleached region at the same time point. These corrected values were then normalized by setting the average pre-bleach intensity to 1 and the intensity of the first frame after photobleaching to zero. We calculated the maximum fractional fluorescence recovery as the average recovery observed for the last three time-points of the time-lapse.

#### Statistical analysis

##### Inter-punctum intervals

**(IPIs)** In this study, we relied on inter-punctum intervals as a standardized measure of the distribution of discrete puncta. This metric has been used previously to analyze the distribution of MEC-4 channels (Arnadottir et al, 2011; Cueva et al, 2007; Vasquez et al, 2014) and here we use it for this purpose and for analyzing the associated BL puncta. Because IPI is derived from the distance between the location of the peak intensity of each punctum and its neighbor, it is more robust to background noise and errors inherent in intensity-based image segmentation. Table S2 collects and summarizes IPI measurements (mean±standard deviation, median ± interquartile range, number of ALMs analyzed, number of IPIs analyzed) as a function of each *C. elegans* strain. For pairwise comparison of cumulative distributions of IPI values, we performed the Kolmogorov-Smirnof test (using the scipy.stats library in Python) and reported the corresponding ks-statistic and p-value on the respective figures. For comparing IPIs between three or more groups, we performed 1-way ANOVA and Tukey-HSD post hoc testing (using the statsmodels library in Python), and reported the results in the figure legend.

##### Other data types

In addition to summarizing IPI measurements, Table S2 also summarizes other measurements derived from still and time-lapse imaging (ICQ measurements, puncta colocalization, MEC-4 fluorescence intensity, maximum fluorescence recovery after photobleaching), *in vivo* whole-cell patch clamp recordings (peak mechanoreceptor currents), and behavioral studies touch sensitivity. We performed Student’s t-test (using the scipy.stats library in Python) for comparison between two groups, 1-way ANOVA with Tukey post hoc testing (using the statsmodels library in Python) for comparison between three or more groups with one independent variable, and 2-way ANOVA with Tukey post hoc testing (using the statsmodels library in Python) for comparison between three or more groups with two independent variables, and reported the results in the figure legend. For determination of statistically significant differences between groups, p-value cutoff was set at 0.05.

## SUPPLEMENTARY INFORMATION TITLES AND LEGENDS

Movie S1: TEM serial section ALML showing discontinuous ECM beams (related to Figure 1E)

Table S1. List of all strains used in screening for localization of proteins along TRNs

Table S2: Summary data tables

**A. Inter-punctum interval**
**B. Punctum length**
**C. ICQ**
**D. Puncta colocalization**
**E. MEC-4 intensity**
**F. FRAP**
**G. Touch score**
**H. Peak MRC amplitude**

***Figure S1. TEM images of TRN cross sections showing discontinuous segments containing beam-like structure in the ECM, related to Figure 1E.***

**A-D**. TEM micrographs showing the cross sections of TRNs at different points along the neuron in a serial section stack. (A) and (B) show the ALMR and AVM neurons, respectively, from the same worm as the ALML shown in Figure 1E. (C) and (D) show the PLM and PVM neurons from a different worm. Red arrows point towards the electron dense ECM beams in the extracellular space between the TRN and the epidermis. z-axis maps of the entire serial section stack of TEM images used for the respective top panels show the relative position of the segments with (red) or without (yellow) visible ECM beams. Red numbers indicate the length (in microns) of each segment with visible ECM beams. Black numbers indicate the distance (in microns) between the mid-points of successive red segments.

**E.** TEM micrographs showing the cross sections of several neurons in the ventral cord, including the PVM (left) or AVM (right). AVM and PVM neurons are highlighted in yellow. Red arrows point towards the electron dense ECM beams in the extracellular space between the TRN and the epidermis. The micrograph of PVM (left) and AVM (right) is from the same series shown in Fig. S1D and Fig. S1B, respectively. Scale bars = 500 nm.

***Figure S2. Punctum length and FRAP fractional recovery for BL proteins and the MEC-4 channel, related to Figure 2.***

**A.** Boxplots showing the distribution of punctum lengths for LAM-1::wSc, LAM-2::mNG, NID-1::wSc and mNG::MEC-4. The dataset used is the same as that in Figure 2B. Box denotes the median and interquartile range, while whiskers extend to the limits of the distribution excluding outliers. One-way ANOVA comparing punctum lengths across different protein labels shows statistically significant differences [*F*(3,11753) = 68.05, *p* = 1.28e-43]. Tukey HSD post hoc analysis shows significant differences (*p* = 0.001, groups connected by gray bars) between all pairs except NID-1::wSc and mNG::MEC-4. Summary data are in Table S2.

**B.** Representative, *in vivo* live imaging of myristoylated GFP (myr::GFP) expressed in the TRNs. Top two panels show fluorescence before and after bleaching, respectively; the bottom panel shows a kymograph (distance-time image) of a 10 s pre-bleach period and 60 s observation period.

**C.** Fluorescence recovery after photobleaching (FRAP) for myr::GFP. About 70% fluorescence recovery for myr::GFP was seen during the observation period. Dark trace and shaded regions show the average and standard deviation of the trajectory of fluorescence recovery for 14 neurons.

**D-E.** Fractional fluorescence recovery at 300 s (D) and 60 s (E) after photobleaching for the different protein labels. The dataset used is the same as in Figure 2D, S2C. One-way ANOVA shows statistically significant differences among groups (Panel D: *F*(3,48) = 68.73, *p* = 2.15e-17; Panel E: *F*(4,61) = 179.11, *p* = 5.64e-33). Tukey HSD post hoc analysis shows significant differences (groups connected by gray bars, *p* = 0.001) between all pairs except NID-1::mNG and LAM-2::mNG.

**F-G.** Plots showing the fraction of green puncta that co-localize with red puncta (F) or vice-versa (G) as a function of genotype. The dataset used is the same as in Figure 2F. Each circle shows a measurement from a single ALM neurite. Black squares show the mean of the filled circles or open circles respectively, and the black bars show standard deviation. In all cases, the fractional puncta colocalization between two fluorescence channels from the same cell (circles) are significantly higher (Student’s t-test, p<0.05) than that measured for simulated pairs (boxplots showing the interquartile (box) and full range (whiskers), excluding outliers). One-way ANOVA showed a significant effect of the protein pair (Panel F: *F*(4,65) = 19.87, *p* = 1.0e-10; Panel G: *F*(4,65) = 11.74, *p* = 3.1e-7). Gray bars connect the groups whose fractional puncta colocalization differ from one another by Tukey post hoc analysis (*p* < 0.005). Summary data are in Table S2.

***Figure S3. Basal lamina proteins in other tissues and TRNs (A-B), co-localization and co-transport of RAB-3, MEC-4 and NID-1 (C-F), and fractional recovery as a function of genotype (G), related to Figure 3.***

**A.** Representative images of animals expressing NID-1::wSc (left) and LAM-2::mNG (right) in control, *mec-5(u444)*, *mec-1(e1738)* and *mec-9(u437)* animals showing the similarity in distribution in basal lamina surrounding the gonad as a function of genotype. Scale bars = 100 µm.

**B.** Fluorescence photomicrographs of animals co-expressing LAM-2::mNG and a cytoplasmic RFP marker in the TRNs revealing discrete LAM-2 puncta associated with the TRNs in control, but not in *mec-5(u444)*, *mec-1(e1738)* and *mec-9(u437)* animals. Scale bars = 10 µm.

**C-E**. Kymographs of dual labeled TRNs showing that mobile mNG::MEC-4 puncta co-localize with mCherry::RAB-3 puncta *in vivo* (C) and in TRNs cultured *in vitro* (D), but do not co-localize with NID-1::wSc puncta *in vivo* (E).

**F.** Plot showing the density (average number of puncta per 20 µm) of all detected mCherry::RAB-3 positive vesicles (magenta) and MEC-4-containing RAB-3 positive vesicles (white) in each cell. The dataset used is the same as in Figure 3D. Pairs of data points from individual cells are connected by magenta lines. Circles denote mean values and thick vertical lines show the standard deviation. One-way ANOVA fails to detect an effect of genotype on total RAB-3 density [*F*(3,80) = 2.22, *p* = 0.09] and on the density of MEC-4-containing RAB-3 puncta [*F*(3,80) = 1.77, *p* = 0.16].

**G.** Plots showing the fractional fluorescence recovery at 300 s after photobleaching for the different genotypes. The dataset used is the same as in Figure 3F. One-way ANOVA shows statistically significant effect of genotype [*F*(2,67) = 3.35, *p* = 0.04] and a post hoc analysis (Tukey HSD) shows a significant difference (groups connected by gray bars) between control and *mec-1(e1738)* but no significant difference between *mec-1* and *mec-9* mutants.

***Figure S4. Mechanosensory ECM assembly is not dependent on the ion-channel subunits or TRN microtubules, related to Figure 4.***

**A.** Representative images of LAM-1::wSc puncta present in *mec-4(u253)mec-10(tm1552)* double mutant (top) and *mec-4(u253)mec-10(tm1552);degt-1(tm6356)* triple mutant (bottom). Anterior is to the left; scale bars are 10 µm.

**B.** Probability density distribution plots of IPIs (top) and the individual IPIs (bottom) for LAM-1::wSc in *mec-4(u253);mec-10(tm1552)* double mutant and *mec-4(u253);mec-10(tm1552)degt-1(tm6356)* triple mutant. White circles denote mean, thin vertical lines denote the median and thick horizontal lines denote the interquartile ranges of the population. Summarized data are collected in Table S2.

**C.** Plots of cumulative probability distribution of LAM-1::wSc IPIs in the indicated alleles (green) vs control worms (gray, replotted from Figure 2B). The results of two-sample Kolmogorov-Smirnov tests are indicated in each plot.

**D.** Stacked bar plots representing the fraction of MEC-4 in mechanosensory complexes (dark green), and in the diffuse pool (light green) in control vs. *mec-7(ok2152)* animals show that MEC-4 is distributed evenly between the two pools in both genotypes. The dataset used is the same as in Figure 4E-F. Student’s t-test shows no significant difference in the fraction of MEC-4 in mechanosensory complexes between the two genotypes (p=0.13). Summarized data are collected in Table S2.

**E-G.** MEC-4 and NID-1 puncta co-localize in control and *mec-7(ok2152)* animals. Plots showing ICQ (E), the fraction of MEC-4 puncta that co-localize with NID-1 puncta (F) or vice-versa (G) as a function of genotype. The dataset used is the same as in Figure 4E-F. Each circle (open circles: control, filled circles: *mec-7(ok2152)*) shows a measurement from a single ALM neurite. Black squares show the mean and the black bars show standard deviation. Observed ICQ and fractional puncta colocalization values for control are significantly higher (gray bars connecting groups with p<0.05 in Student’s t-test) than *mec-7(ok2152)* in E, F and G. For both genotypes, the ICQ (E) or fractional puncta colocalization values (F,G) between two fluorescence channels from the same cell (circles) are significantly higher (p-values<0.05 in Student’s t-test) than that measured for simulated pairs (boxplots, box: interquartile range, whiskers: full range excluding outliers). Summarized data are collected in Table S2.

***Figure S5. Effect of loss of epi-1 and nid-1 on MEC-4 puncta (A-D), MEC-4 co-localization with BL proteins (E-G), and voltage-gated currents in ALM as function of genotype (H-I), related to Figure 5.***

**A.** Representative images showing mNG::MEC-4 localize to discrete puncta distributed along the ALM neurons and TRN morphology defects in *epi-1(gm57)* adult animals. Boxed area in the top panel is expanded in the lower panel. Similar patterns observed in 9 ALM neurons in 9 adult animals.

**B.** Probability density distribution plots of IPIs (top) and the individual IPIs (bottom) for mNG::MEC-4 in *epi-1(gm57)* mutant animals. Due to the TRN morphological defects the complete TRNs could not be analyzed in their entirety; instead IPIs were measured in 13 segments that were at least 50 µm long drawn from 9 independent biological replicates (ALM neurites). White circles denote mean, thin vertical lines denote the median and thick horizontal lines denote the interquartile ranges of the population. Summarized data are collected in Table S2.

**C.** Cumulative distribution plots of mNG::MEC-4 puncta distribution in *epi-1(gm57)* mutants and control animals. The green and gray lines are data for *epi-1(gm57)* and control, respectively. Control IPIs were computationally selected from 20 randomly positioned 50 µm ALM segments in 15 control animals to provide a better match to the analysis available for *epi-1(gm57)* animals. The results of two-sample Kolmogorov-Smirnov tests showing no significant difference between the two distributions are indicated in the plot.

**D.** Stacked barplots representing the fraction of MEC-4 in mechanosensory complexes (dark green), and in the diffuse pool (light green) in control vs. *nid-1(cg119)* mutant animals show that *nid-1* loss of function redistributes MEC-4 to a diffuse membrane pool. The dataset used is the same as in Figure 5E-F. Numbers in the bar graph indicate the sample size (number of ALMs) and error bars show standard deviations of the measurements for each genotype. Student’s t-test shows a significant difference (gray bar) in the MEC-4 diffuse pool between the genotypes (p=2.28e-21). Summarized data are collected in Table S2.

**E-G.** Loss of *nid-1* function decreases MEC-4 and LAM-1 intensity correlation, even though individual puncta co-occur more often than expected from a random distribution. Plots showing ICQ (E), the fraction of MEC-4 puncta that co-localize with LAM-1 puncta (F) or vice-versa (G) as a function of genotype. The dataset used is the same as in Figure 5E-F. Each circle (open circles: control, filled circles: *nid-1(cg119)*) shows a measurement from a single ALM neurite. Black squares show the mean of the filled circles or open circles respectively, and the black bars show standard deviation. Observed ICQ and fractional puncta colocalization values for control are significantly higher (Student’s t-test, p<0.05) than *nid-1(cg119)* in E, F and G. For both genotypes the ICQ (E) or fractional puncta colocalization values (F,G) between two fluorescence channels from the same cell (circles) are significantly higher (gray bars connecting groups with p-values<0.05 in Student’s t-test) than that measured for simulated pairs (boxplots, box: inter-quartile range, whiskers: full range excluding outliers). Summarized data are collected in Table S2.

**H.** Representative membrane current traces evoked by voltage pulses (from ™80 to +80mV, in 20 mV increments) applied from a holding potential of ™60 mV. Each trace represents the average of three stimulus presentations for control (left, dark gray), *nid-1(cg119)* (middle, green) and *mec-5(u444)* (right, magenta) mutants. Traces are shaded according to the voltage applied during the step. Similar results obtained in a total of 11, 9 and 2 recordings for control, *nid-1* and *mec-5* animals, respectively.

**I.** Peak current vs. voltage. Dark gray, dark green and magenta traces are the average peak current of the lighter gray, green and magenta traces for control, *nid-1* and *mec-5* mutants, respectively. For individual recordings, membrane voltage is corrected for residual series resistance.

***Figure S6. NID-1[ΔG3], LAM-2[NSS] and LAM-3[NSS] mutations successfully disrupt nidogen/laminin binding, related to Figure 6.***

**A.** Stacked bars representing the fraction of MEC-4 in the diffuse pool (light green) and mechanosensory complexes (dark green) as a function of genotype. One-way ANOVA shows a significant effect of genotype [*F*(6,188) = 148.43, *p* = 1.51e-68]. Gray bars connect groups that are significantly different (p<0.05) from control by Tukey-HSD post hoc analysis. Numbers on bars indicate the number of biological replicates for each genotype. Summary data in Table S2.

**B.** Touch response as a function of genotype. One-way ANOVA shows significant differences between groups [*F*(6,518) = 29.88, *p* = 8.51e-31]. gray bars connect groups which are significantly different (p<0.05) from control by Tukey-HSD post hoc analysis. Bars are the average (± SEM) response to 10 touches of *n=75* animals tested blind to genotype in 3 independent cohorts. Summarized data are collected in Table S2.

**C.** Summary of the NID-1 localization in TRNs, pharynx and body wall muscles in the different genotypes. “+++” indicates strong localization, “+” indicates weak localization and “-” indicates no localization. NID-1 localization data was not collected for the NID-1[ΔG1-G2] background. Loss of NID-1 protein in the *nid-1(cg119)* null background was confirmed by Western blot in previous literature (Kang and Kramer, 2000).

**D-E.** Representative images showing localization of nidogen (NID-1, magenta) and laminin-γ (LAM-2, green) to BL surrounding the pharynx (D, white arrows), nerve ring (D, white arrowheads), body wall muscle (E, black arrows) and TRN (E, black arrowheads) in the indicated mutants. Nidogen fluorescence around the pharynx and body wall muscles is lost in NID-1[ΔG3] and LAM-2[NSS]LAM-3[NSS] backgrounds. In the LAM-2[NSS] single mutant background, nidogen is partially lost from the basal lamina surrounding the pharynx and body wall muscles, indicating LAM-3 is capable of recruiting nidogen without LAM-2. In the LAM-3[NSS] single mutant, nidogen was recruited to all tissues similar to control, as LAM-2 is present in all laminin and LAM-2 is capable of recruiting nidogen without LAM-3. Nidogen fluorescence in the nervous tissue-associated BL, such as the nerve ring and puncta along TRNs are preserved in all the mutants tested, indicating the role of additional factors that recruit nidogen to these BL. In contrast, localization of laminin to the pharynx and body wall muscle is not affected in any of the mutants disrupting nidogen-laminin binding.

***Figure S7. Disrupting Ku15 in MEC-1 minimally affects LAM-1, LAM-2 and NID-1 puncta distribution, related to Figure 7.***

**A-B**. Representative images of ALM neurons in dual-color transgenic animals showing preservation of ECM puncta (LAM-1 in A, NID-1 and LAM-2 in B) but loss of MEC-4 (A) puncta in the *mec-1(e1526)* allele. Anterior is to the left, scale bar is 10 µm.

**C.** Probability density distribution plots of IPIs (top) and the individual IPIs (bottom) in *mec-1(e1526)* mutants for LAM-1::wSc and mNG::MEC-4, NID-1::wSc (E) and LAM-2::mNG (E). White circles denote mean, thin vertical lines denote the median and thick horizontal lines denote the interquartile ranges of the population. Summarized data are collected in Table S2.

**D.** Plots of cumulative probability distribution of LAM-1::wSc (top) and mNG::MEC-4 (bottom) IPIs in the *mec-1(e1526)* mutant (magenta or green) vs control worms (gray, replotted from Figure 5F). The results of two-sample Kolmogorov-Smirnov tests are indicated in each plot.

**E.** Probability density distribution plots of IPIs (top) and the individual IPIs (bottom) in *mec-1(e1526)* mutants for LAM-1::wSc and LAM-2::mNG. White circles denote mean, thin vertical lines denote the median and thick horizontal lines denote the interquartile ranges of the population. Summarized data are collected in Table S2.

**F.** Plots of cumulative probability distribution of NID-1::wSc (top) and LAM-2::mNG (bottom) IPIs in the *mec-1(e1526)* mutant (magenta or green) vs control worms (gray). The results of two-sample Kolmogorov-Smirnov tests are indicated in each plot.

**G-I.** Residual MEC-4 puncta colocalize with LAM-1 puncta in *mec-1(e1526)* animals. Plots showing ICQ of LAM-1 and MEC-4 (G), the fraction of MEC-4 puncta that co-localize with LAM-1 puncta (H) or vice-versa (I) as a function of genotype. Each circle (open circles: control, filled circles: *mec-1(e1526)*) shows a measurement from a single ALM neurite. Black squares show the mean of the filled circles or open circles respectively, and the black bars show standard deviation. Observed values for control are significantly higher (Student’s t-test, p<0.05) than *mec-1(e1526)* in G-I (gray bars). For both genotypes the ICQ (G) or fractional puncta colocalization values (H,I) between two fluorescence channels from the same cell (circles) are significantly higher (Student’s t-test, p<0.05) than that measured for simulated pairs (boxplots, box: inter-quartile range, whiskers: full range excluding outliers). Summarized data are collected in Table S2.

**J-L.** LAM-2 and NID-1 puncta co-localize in control and *mec-1(e1526)* animals. Plots showing ICQ of NID-1 and LAM-2 (J), fraction of LAM-2 puncta that co-localize with NID-1 puncta (K) or vice-versa (L) as a function of genotype. Each circle (open circles: control, filled circles: *mec-1(e1526)*) shows a measurement from a single ALM neurite. Black squares show the mean of the filled circles or open circles respectively, and the black bars show standard deviation. ICQ values for control are not significantly different (Student’s t-test, p=0.09) from *mec-1(e1526)* (J). There are small but statistically significant differences in fractional puncta colocalization values (K, p=0.006; L, p=0.049) between control and *mec-1(e1526)* animals (gray bars). For both genotypes the ICQ (J) or fractional puncta colocalization values (K,L) between two fluorescence channels from the same cell (circles) are significantly higher (Student’s t-test, p<0.05) than that measured from simulated pairs (boxplots, box: inter-quartile range, whiskers: full range excluding outliers). Summarized data are collected in Table S2.

