## Supplementary figures and images for "Conserved basal lamina proteins, laminin and nidogen, are repurposed to organize mechanosensory complexes responsible for touch sensation"

### Supplemental Figures S1-S7

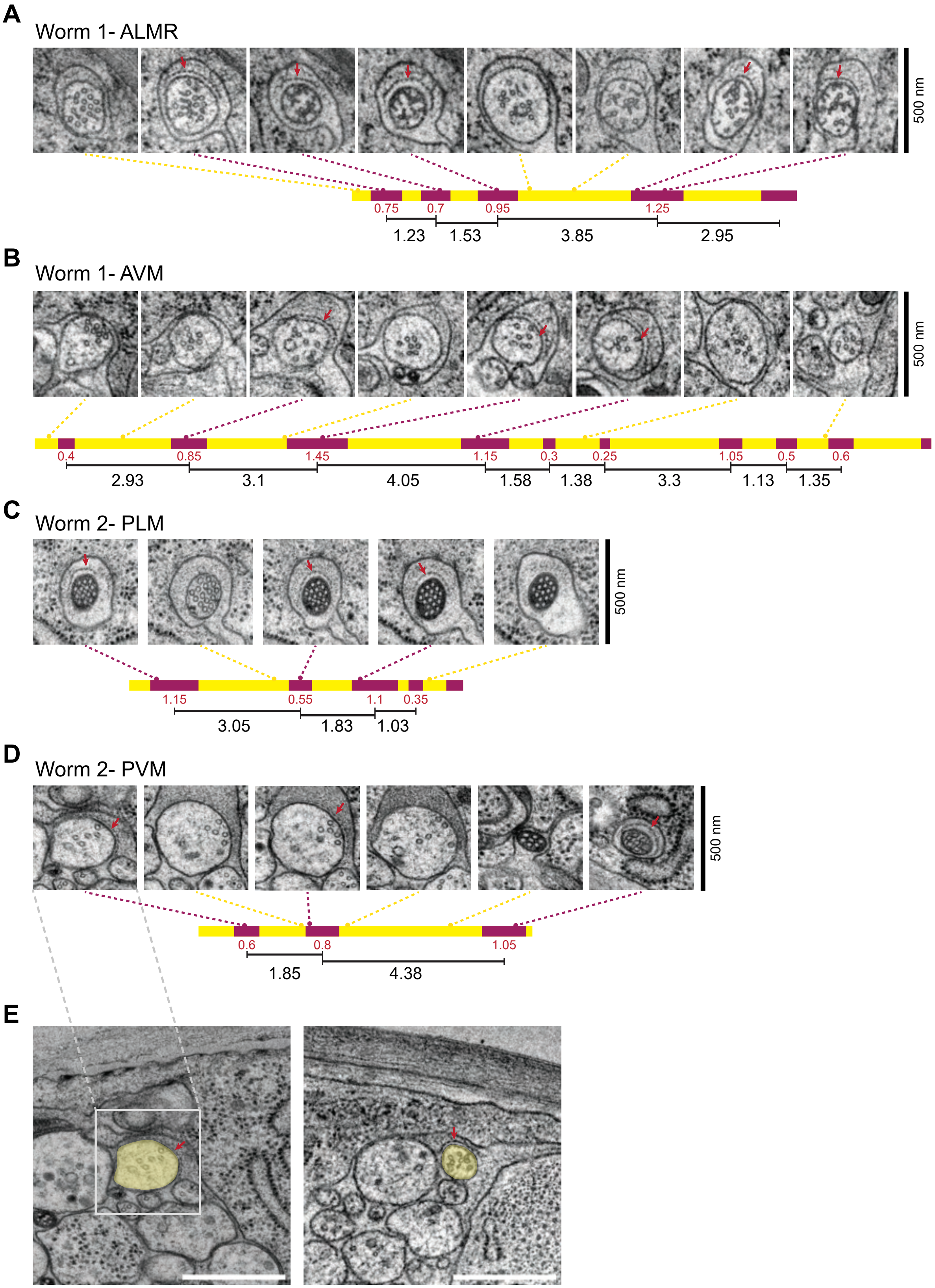

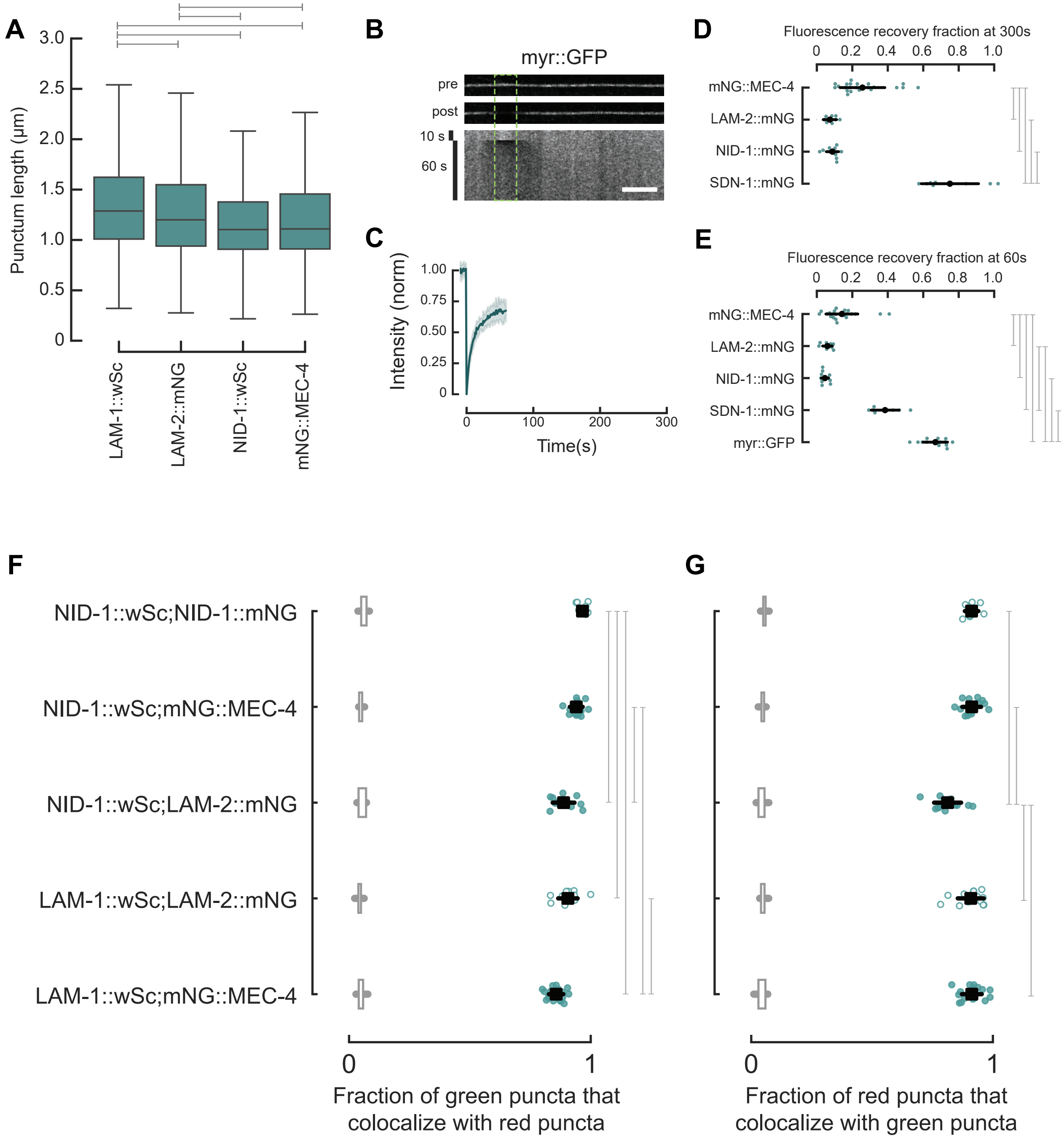

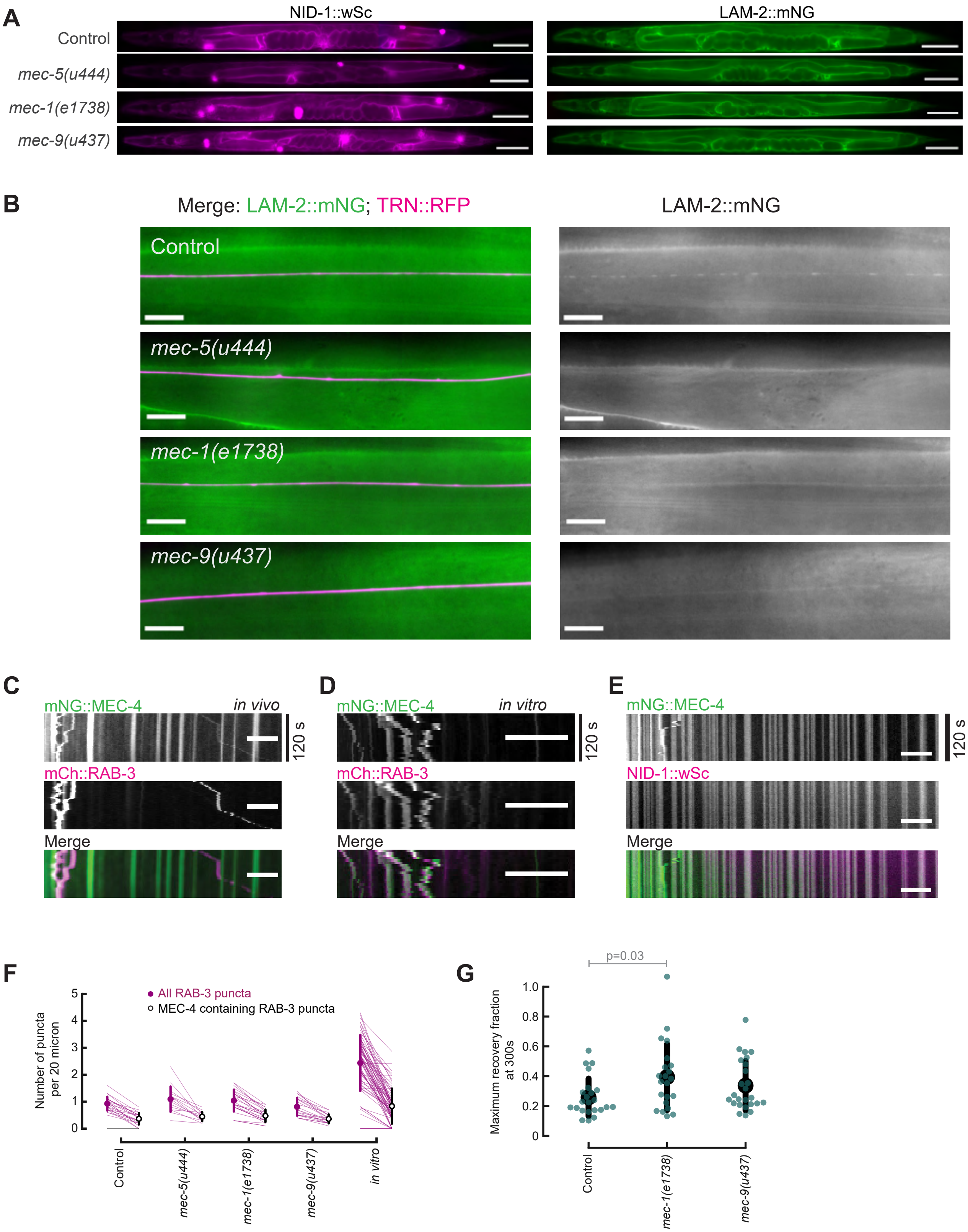

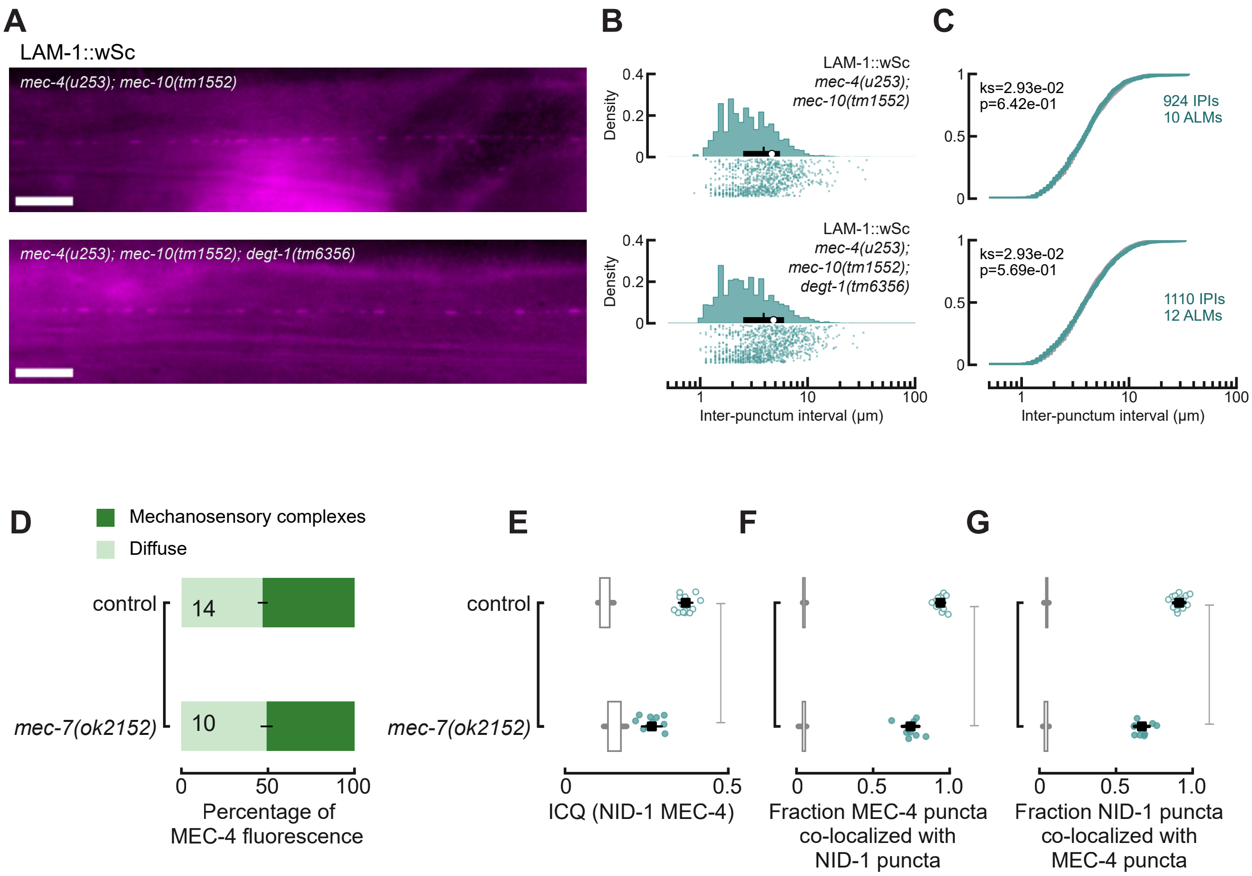

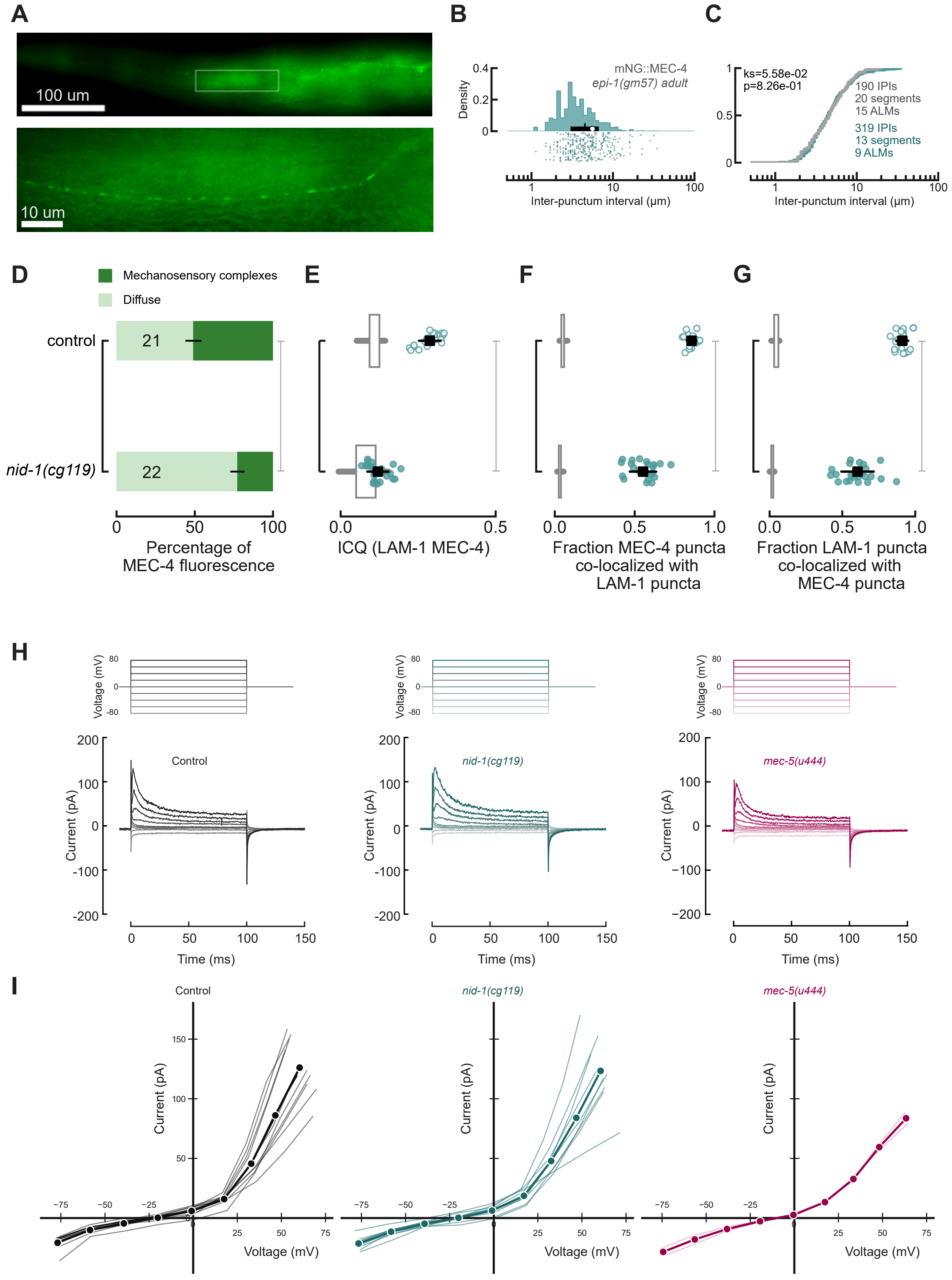

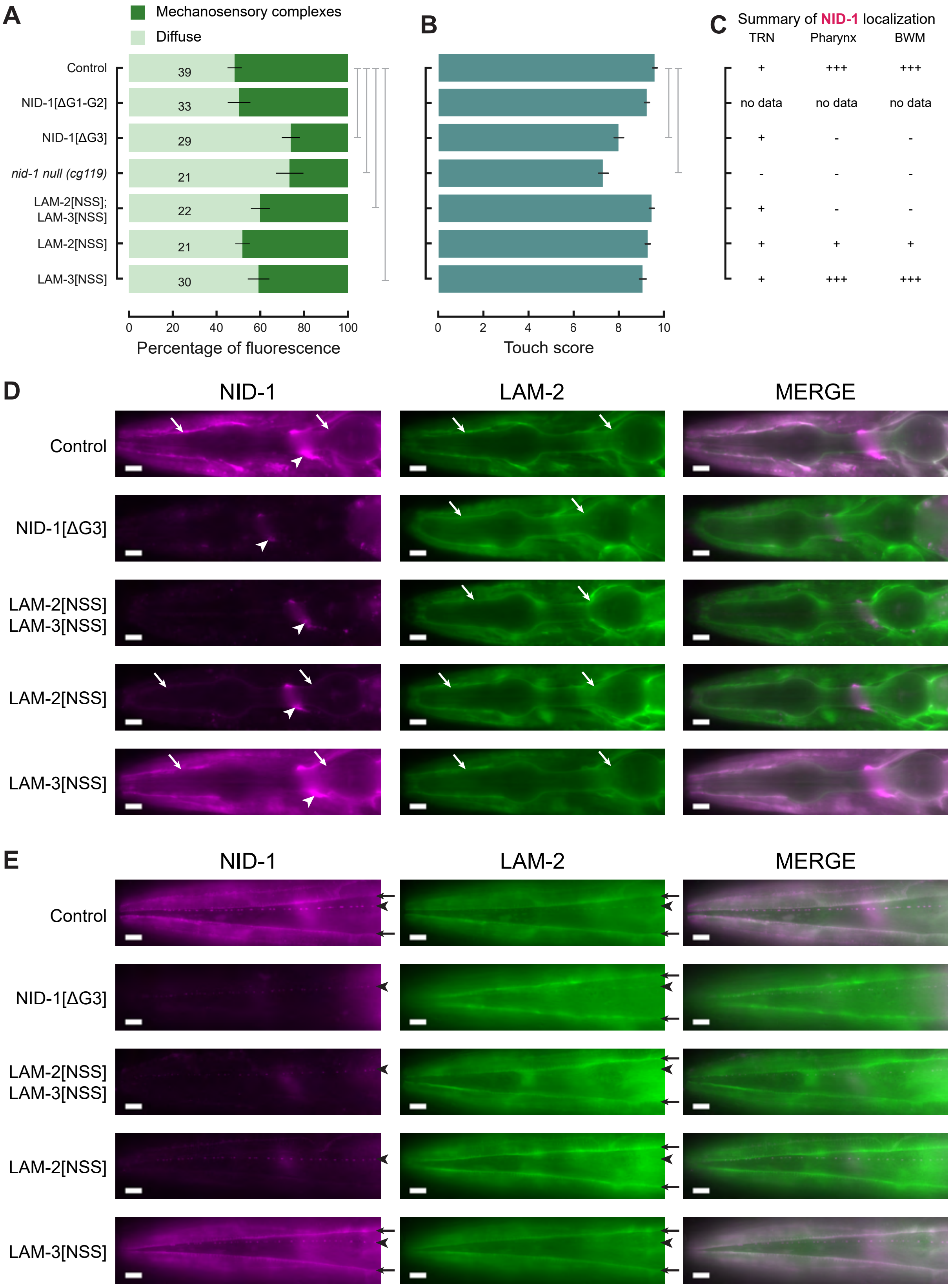

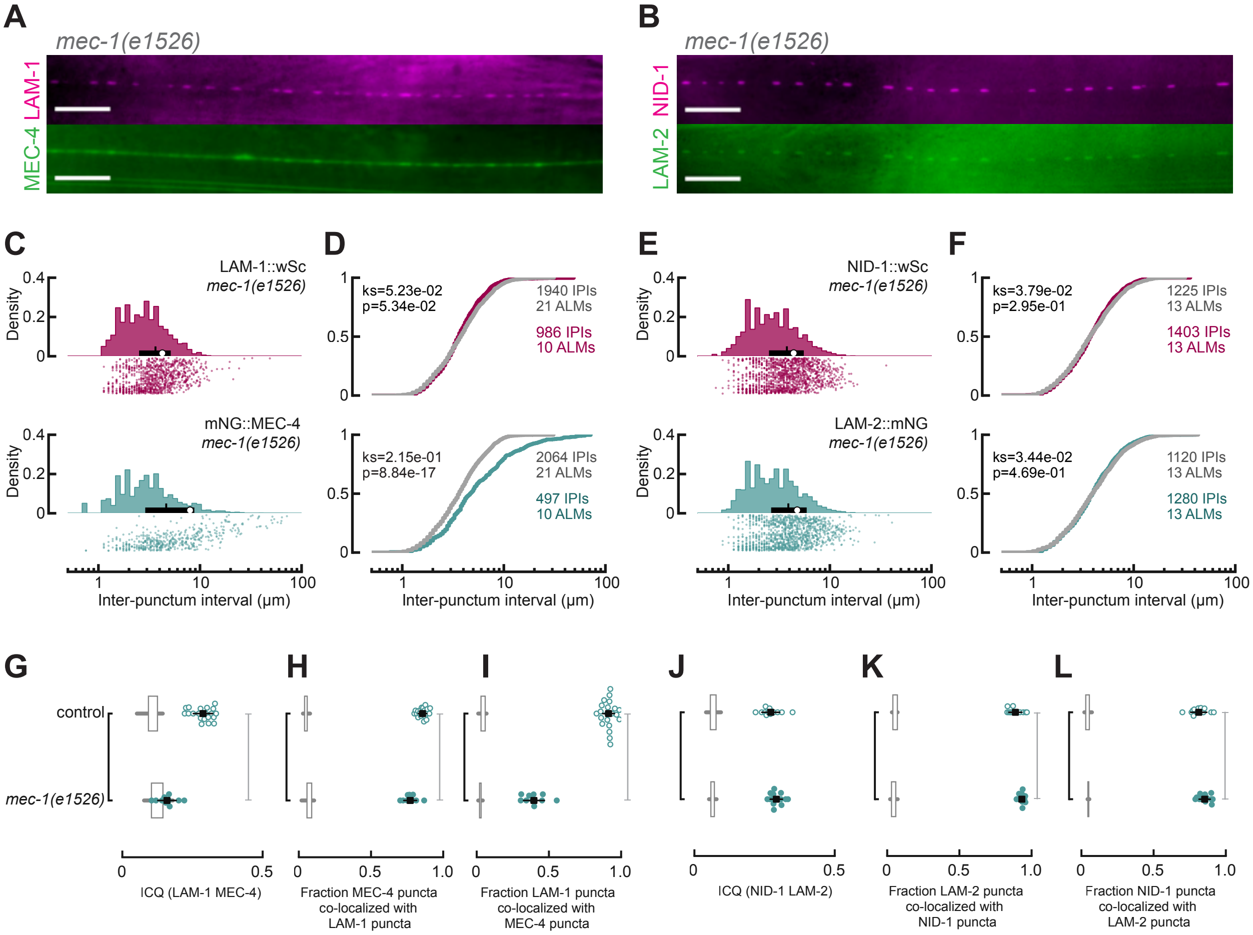
