## Supplemental Table S1 for "Conserved basal lamina proteins, laminin and nidogen, are repurposed to organize mechanosensory complexes responsible for touch sensation"

### Table S1: *List of all strains used in screening for localization of proteins along TRNs*

| Strain name | Genotype | Fluorescence visible around TRN (Yes/No) | If yes, description | Mean touch response for 10 trials (Mean±SEM) | Sample size for touch test (#worms) |
|---|---|---|---|---|---|
| TH502 | *unc-119(ed3) III; ddIs290[sax-7::TY1::EGFP::3xFLAG(92C12) + unc-119(+)].* | Yes | Uniform | 9.81±0.08 | 75 |
| ML2501 | *let-805(mc73[let-805::gfp + unc-119(+)]) unc-119(ed3) III* | Yes | Uniform | 9.79±0.07 | 75 |
| NK2442 | *mig-6(qy37[mNG+loxP::mig-6]) V* | Yes | Uniform | 9.85±0.06 | 75 |
| NK2479 | *pat-2(qy49[pat-2::2xmNG]) III* | Yes | Uniform | 9.61±0.09 | 75 |
| GN800 | *pat-2(pg125[pat-2::lox2272::wSc::3XFLAG]) III* | Yes | Uniform | N.D. | |
| NK2413 | *sdn-1(qy29[sdn-1::mNG+loxP]) X.* | Yes | Uniform | 9.72±0.08 | 75 |
| NK2422 | *him-4(qy33[him-4::mNG+loxP]) IV.* | Yes | Uniform | 9.6±0.12 | 75 |
| NK2555 | *unc-52(qy75[mNG+loxP::unc-52]) II* | Yes (very dim) | Uniform | 8.87±0.22 | 75 |
| NK2583 | *unc-52(qy80[mNG+loxP (synthetic exon)::unc-52]) II* | Only proximal PLM | Patchy | N.D. | |
| NK2500 | *unc-52(qy53[unc-52::mNG+loxP]) II* | No | | N.D. | |
| NK2579 | *fbl-1(qy62[mNG+loxP::fbl-1]) IV* | Yes | Uniform | 1.67±0.2 | 75 |
| NK2404 | *epi-1(qy31[epi-1::mNG+loxP]) IV.* | Yes (very dim) | Uniform | 9.51±0.12 | 75 |
| GN1060 | *lam-1(pg136[lam-1::wSc::Lox2272::3xMyc])/+ IV; uIs31 III* | Yes | Punctate | 9.63±0.09 | 75 |
| NK2335 | *lam-2(qy20[lam-2::mNG+LoxP]) X* | Yes | Punctate | 9.45±0.1 | 75 |
| NK2443 | *nid-1(qy38[nid-1::mNG+loxP]) V.* | Yes | Punctate | 9.59±0.08 | 74 |
| GN973 | *nid-1(pg138[nid-1::wSc::Lox2272::3xMyc]) V; uIs31 III* | Yes | Punctate | 9.32±0.12 | 75 |
| NK2477 | *ptp-3(qy47[ptp-3::mNG+loxP]) II* | Maybe | Patchy | N.D. | |
| NK2353 | *agr-1 (qy27[agr-1::mNG+loxP]) II* | No | | N.D. | |
| NK2456 | *ddr-1(qy43[ddr-1::mNG+loxP]) X* | No | | N.D. | |
| NK2457 | *ddr-2(qy44[ddr-2::mNG+loxP]) X* | No | | N.D. | |
| NK2318 | *dgn-1(qy18[dgn-1::mNG]) X* | No | | N.D. | |
| NK2604 | *emb-9 (qy89[emb-9::mEos2+loxP]) III* | No | | N.D. | |
| NK2326 | *emb-9(qy24[emb-9::mNG +loxP]) III* | No | | N.D. | |
| GN947 | *epi-1(pg139[epi-1::wSc::Lox2272::3xMyc]) IV* | No | | N.D. | |
| NK2590 | *gon-1(qy45[gon-1::mNG+loxP]) IV* | No | | N.D. | |
| NK2581 | *gpn-1(qy35[gpn-1::mNG+loxP]) X* | No | | N.D. | |
| LP172 | *hmr-1(cp21[hmr-1::GFP + LoxP]) I* | No | | N.D. | |
| NK2476 | *ina-1(qy46[ina-1::mKate+loxP]) III* | No | | N.D. | |
| NK2425 | *lam-3(qy28[lam-3::mNG+loxP]) I.* | No | | N.D. | |
| GN956 | *lam-3(pg140[lam-3::wSc::Lox2272::3xMyc]) I* | No | | N.D. | |
| NK2558 | *let-2(qy61[let-2::mNG+loxP]) X* | No | | N.D. | |
| NK2582 | *lon-2(qy55[lon-2::mNG+loxP]) X* | No | | N.D. | |
| FDU1056 | *mig-17(shc19[mig-17::mNG +LoxP])* | No | | N.D. | |

| | | | | |
|---|---|---|---|---|
| NK2557 | *mig-6(qy73[mig-6::mNG+loxP]) V* | No | | N.D. |
| NK2502 | *ten-1(qy56[ten-1::mNG+loxP]) III* | No | | N.D. |
